# Local adaptation of a dominant coastal tree to freshwater availability and solar radiation suggested by genomic and ecophysiological approaches

**DOI:** 10.1101/378612

**Authors:** Mariana Vargas Cruz, Gustavo Maruyama Mori, Caroline Signori Müller, Carla Cristina da Silva, Dong-Ha Oh, Maheshi Dassanayake, Maria Imaculada Zucchi, Rafael Silva Oliveira, Anete Pereira de Souza

## Abstract

Local adaptation is often a product of environmental variations in the geographical space and has implications for biodiversity conservation. We investigated the role of latitudinal heterogeneity in climate on the organization of genetic and phenotypic variation in the dominant coastal tree, *Avicennia schaueriana*. In a common garden experiment, samples from an equatorial region, marked by rainy/dry seasons, accumulated less biomass, showed lower stomatal conductance and transpiration, narrower xylem vessels, smaller leaves and higher reflectance of long wavelengths (red light) on the stem epidermis, than samples from a subtropical region, marked by warm/cold seasons. Transcriptome differences identified between trees sampled under field conditions at equatorial and subtropical sites, were enriched in functional categories as responses to temperature, solar radiation, water deficit, photosynthesis and cell wall biosynthesis. The diversity based on thousands of SNP loci revealed a north-south genetic structure. Remarkably, signatures of selection were identified in loci associated with photosynthesis, anthocyanin accumulation and the responses to osmotic and hypoxia stresses. Our results suggest the existence of divergence in key resource-use characteristics, likely driven by climate seasonality, based on water-deficit and solar radiation. These findings provide a basis for conservation plans and for predictions for coastal plant responses to climate change.

## Introduction

Adaptation is frequently a consequence of spatial variations in selective environmental forces acting on phenotypic diversity^1, 2^. As selective forces operate, they may reduce the heritable variation within a population, leading to the specialization of individuals^2^. Conversely, in highly stochastic environments, selection can increase the species potential for phenotypic plasticity^3^. Therefore, the investigation of adaptation is fundamental for understanding the ability of a species to respond to environmental changes^4, 5^.

Alarmingly high rates of environmental changes have been observed over the last decades, particularly affecting coastal ecosystems^6–9^, for instance, bleaching of coral reefs worldwide^10^, large-scale dieback of mangrove communities^11^ and increased mortality of seagrass beds^12^. Further environmental changes and coastal biodiversity loss threaten critical socioeconomic and ecological services provided by coastal ecosystems, including their pivotal role as blue carbon stocks^13^. In this context, studies on the organization of adaptive variation of coastal habitat-forming species are necessary to improve predictions on the impacts of future climate conditions on marine ecosystems^14^ and to support efficient conservation practices^15^.

Coastal habitat-forming species are frequently water-dispersed over long distances^16, 17^ and often occur across wide latitudinal ranges. The connectivity among populations of these species may be influenced, not only by their dispersion capabilities but also by means of natural selection, caused by varying establishing success over the broad environmental heterogeneity across latitudes^18^. For instance, distinct coastal species in the North Atlantic show a north-south organisation of diversity with evidences of selection for different thermal regimes^19–21^. Similarly, in the Southwest Atlantic, an overlapping north-south structure of the genetic diversity, with reduced gene flow between northern and southern populations, has been observed in phylogenetically distant habitat-forming species^22–24^. In this context, one could expect northerly and southerly populations of these species to adapt differently to contrasting environments over their latitudinal ranges, especially due to their large population sizes^2, 25^. However, the neutrality of the molecular markers used in previous studies has precluded inferences regarding adaptive variation. The existence of local adaptation remains virtually unknown in species that play a central role in sustaining the coastal biodiversity in the South Atlantic. This limited knowledge about the organization of non-neutral variation compromises accurate predictions and suitable conservation efforts for sustainable management.

Here, we tested the hypothesis that latitudinal heterogeneity in climate variables shape the variation of genotypes and phenotypes involved in the optimisation of resource-use in widely distributed coastal species. To reduce the potential for incorrect conclusions about selection^26^, we integrated three independent, but complementary approaches, making predictions as follows: (1) using a common garden experiment, individuals from contrasting latitudes would show genetically-based phenotypic divergence in ecophysiological traits; (2) under contrasting latitudes, transcriptomic changes would be detected in genes involved in responses to environmental variation; and (3) signatures of selection along the species distribution would be detected in genes involved in responses to latitudinal variation in air temperature, solar radiation and freshwater availability determined by air water vapour pressure deficit (VPD), rainfall and tidal regime. We considered the new-world coastal tree species, *Avicennia schaueriana* Stapf & Leechman ex Moldenke, found in the Lower Lesser Antilles and widespread from Venezuela to the southernmost temperate mangroves on the Atlantic coast of South America (∼28 °S)^27^ as a model to test this hypothesis. The broad latitudinal range of *A. schaueriana*, spanning wide environmental variation (Fig. 1), and the previously detected north-south structure of neutral variation^22^, facilitate ecological predictions and the accumulation of divergence, motivating this choice. Our results indicated that fluctuations in VPD, rainfall and solar radiation may be associated with the observed phenotypic and genotypic variation as well as the regulation of gene expression in this dominant coastal tree. We discuss the implications of our results for the persistence of coastal biodiversity in the context of climate change.

**Figure 1.**
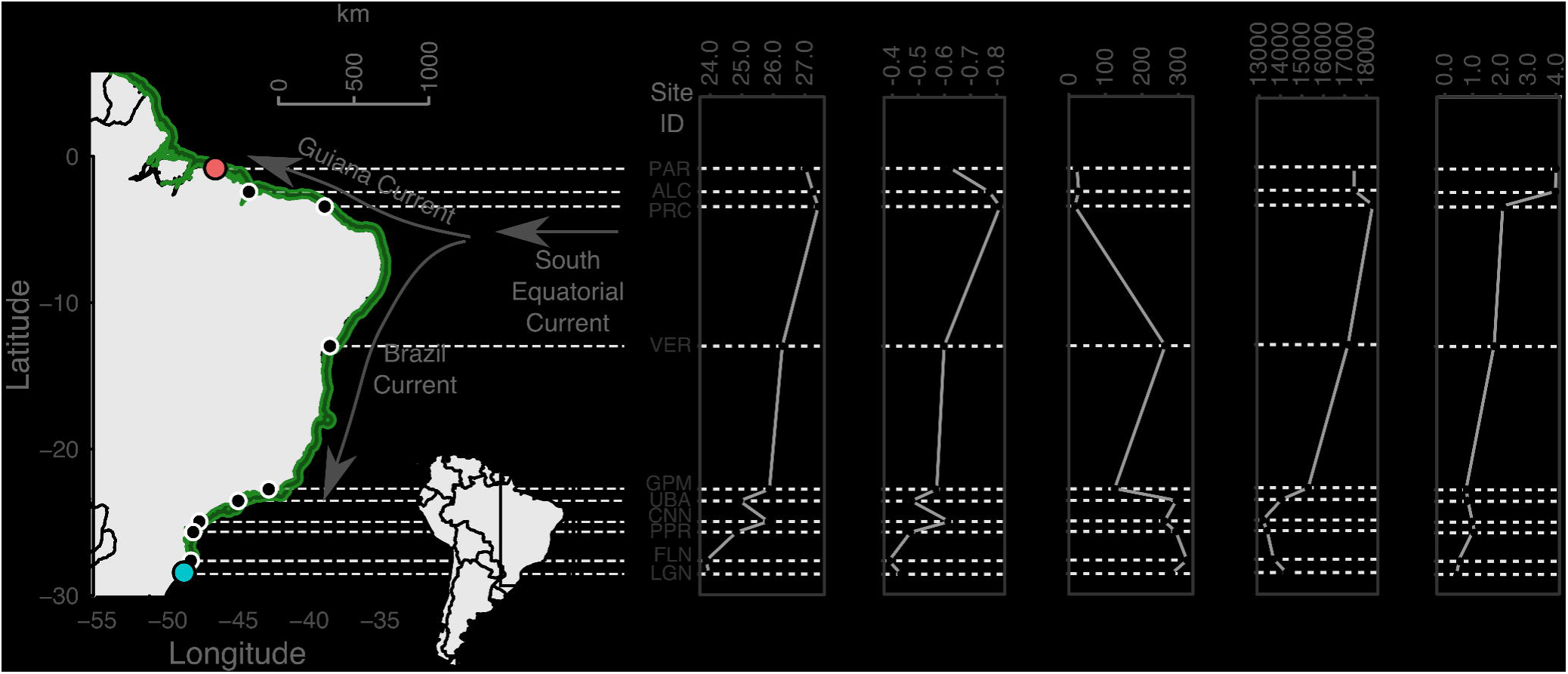
Geographical locations of *Avicennia schaueriana* sampling sites. Characterisation of *Avicennia schaueriana* sampling sites along a latitudinal gradient on the Atlantic coast of South America. The green-shaded area represents the presence of the species; coloured points represent sampling sites of propagules, used in a common garden experiment, and of plant tissues, used for both DNA- and RNA-sequencing; black points represent sampling sites of plant tissues, used for genomic DNA-sequencing only; arrows represent the flow directions of major oceanic currents. Environmental variation across sampling sites was obtained from public databases of climate (‘WorldClim’^111, 112^) and tide variables (‘Environment Climate Data Sweden’^110^).

## Materials and Methods

### Propagule sampling

Mature propagules were collected from 15 *Avicennia schaueriana* Stapf & Leechman ex Moldenke mother trees that were at least 100 m apart from one another, at the following two genetically contrasting populations^22^: (1) the southernmost range limit of American mangroves, in the subtropical region, and (2) an equatorial site in one of the world’s largest macrotidal mangrove forests^28, 29^, near the northernmost limit of the species range (Fig. 1). We refer to samples collected in the former site as “subtropical” and those in the latter as “equatorial” throughout this work. Sampling in more localities was impracticable due to the absence of mature propagules during fieldwork. A detailed characterisation of each of these contrasting sites can be found in Table 1, in the Supplementary Methods and in the Supplementary information Figure S1.

**Table 1.**
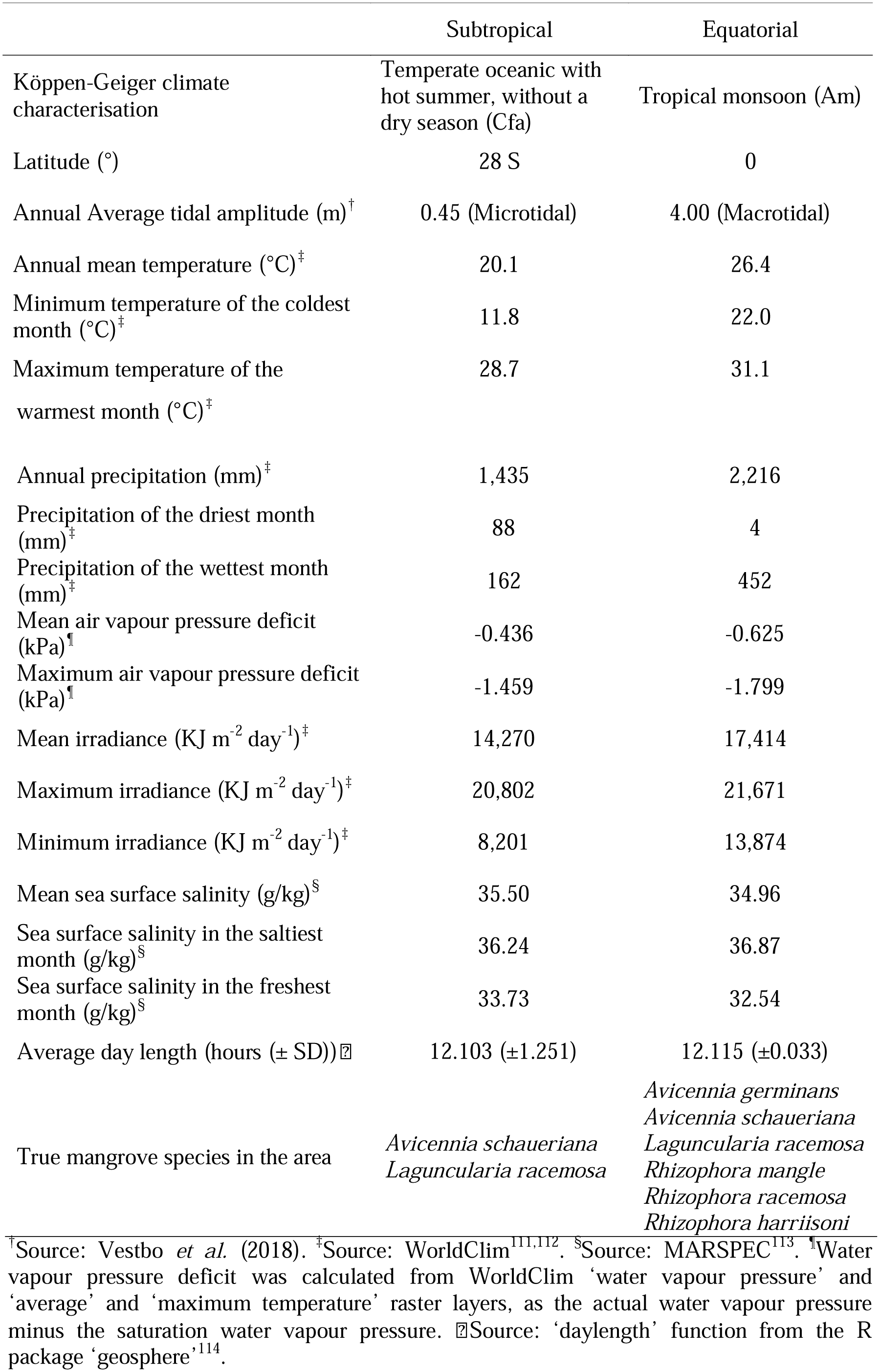
Characterisation of subtropical and equatorial sampling sites of propagules used in the common garden experiment and of plant material used for RNA-sequencing. SD: standard deviation.

### Comparative ecophysiology using a common garden experiment

Propagules were germinated as described for *Avicennia germinans*^30^. We grew propagules in trays with local mangrove soil for two months. After this period, 44 similar-sized seedlings from 30 distinct mother trees, 15 from equatorial and 15 from subtropical sites — with an average height of 18 cm, most with three leaf pairs and senesced cotyledons — were transplanted to 6 L pots filled with topsoil and sand (1:1). Seedlings were cultivated for seven months under homogenous conditions in a glasshouse at the University of Campinas, São Paulo, Brazil (22°49’ S 47°04’ W), where automatic sensors coupled with a data logger (Onset Computer Corp.) measured the atmospheric humidity and temperature every 15 minutes. Seedlings were irrigated daily at 10 a.m. and 5 p.m. with a 3-minute freshwater spray. Twice a week, nutrients were added to the soil using 600 mL of 0.4X Hoagland solution with 15.0 g L^-1^ NaCl per pot. Pots were rotated weekly to reduce the effects of environmental heterogeneity. The environmental conditions in the glasshouse were different from those at each sampling site, as is shown in the Supplementary information Fig. S2.

The light reflectance of stems was measured in ten plants from each sampling site using a USB4000 spectrophotometer (OceanOptics, Inc.) coupled to a deuterium-halogen light source (DH-2000; OceanOptics) using a light emission range from 200-900 nm. Photosynthesis, stomatal conductance and transpiration rates were measured every 2.0-2.5 hours in five six-month-old individuals from each sampling site on two different days using a Li-Cor 6400 XT (Li-Cor Corp.).

After harvest, three plants without flowers or flower buds from each sampling site were split into leaves, stems and roots, washed with distilled water, dried for 7 days at 70 °C and weighed. The individual leaf area, total leaf area and leaf lamina angle per plant were measured through photographic analyses using ImageJ^31^. The specific leaf area (SLA, cm^2^ leaf area kg^-1^ leaf dry biomass) was also calculated for these samples. Stems were fixed in FAA (Formaldehyde Alcohol Acetic acid), stored in 70% alcohol for wood anatomy analysis and cut into 30-μm thick transverse sections. Sections were stained with a mixture of 1% Astra Blue and 50% alcohol (1:1) followed by 1% Safranin O. Micrographs were taken using an Olympus BX51 microscope coupled to an Olympus DP71 camera (Olympus Corp.). The following wood traits were quantified using ImageJ and R v.4.0.0: vessel lumen area (A), vessel density in xylem (number of vessels/xylem area), vessel grouping index (mean number of vessels per vessel grouping) and vessel lumen area in sapwood (vessel lumen area/sapwood area). The vessel arithmetic diameter (D) was estimated according to Scholz et al.^32^.

Shapiro-Wilk’s normality tests and Fisher’s F-tests of equality of variances were performed in R v.4.0.0 to determine the suitability of group comparison statistical tests. In comparisons between equatorial and subtropical samples, we used the unpaired Student’s T-tests or, alternatively, Mann-Whitney-Wilcoxon tests. Multiple-group comparisons were conducted for the analysis of environmental conditions in the field vs. in the glasshouse using one-way analysis of variance (ANOVA) with Tukey’s post hoc honest significant difference (HSD) tests. A significance level of 0.05 was used as the alpha for all tests. The Bonferroni adjustment of P-values was conducted for multiple comparisons.

### Plant material for RNA extraction and RNA-sequencing

Plant material used for RNA-sequencing (RNA-Seq) was collected from the equatorial and subtropical sites described in the “Propagule sampling” section and immediately stored in RNAlater (Ambion, Inc.). Because precise species identification requires the analysis of both vegetative branches and flower morphology, opportunities of sampling were limited by the reproductive phenology at each visited site. Leaves, stems and flowers from three adult trees at least 100 m apart were collected from July-August of 2014, which corresponds to the end of winter at the subtropical site and the beginning of the dry season at the equatorial site. To minimize the detection of differential transcripts expression due to circadian changes, sampling was conducted during low tide and from 10:30 AM to 4:00 PM. A detailed description of the environmental conditions at the time of sampling is available in the Supplementary information Table S1.

We extracted RNA according to Oliveira et al.^33^ and evaluated its integrity and purity using agarose gel electrophoresis and a NanoVue spectrophotometer (GE Healthcare Life Sciences). lllumina TruSeq RNA Sample Preparation kits (Illumina, Inc.) were used to construct cDNA libraries. cDNA quality was assessed using the Agilent 2100 Bioanalyzer (Agilent Technologies) and concentrations were quantified by qPCR using the Sequencing Library qPCR Quantification kit (Illumina, Inc.). Sequencing was performed using two 36-cycle TruSeq SBS paired-end kits (Illumina, Inc.) and the Genome Analyzer IIx platform (Illumina, Inc.).

### Assembly and characterisation of the <u>A. schaueriana</u> transcriptome

Adapter sequences were trimmed, and 72-bp paired-end reads were filtered by quality (Phred score ≥ 20 for at least 70% of the read length) using the NGS QC Toolkit v.2.3^34^. High-quality reads were used for transcriptome assembly in the CLC Genomics Workbench (https://www.qiagenbioinformatics.com/). We used the default settings, except for the distance between pairs (300-500 bp) and k-mer size (45 bp).

Reads were mapped to transcripts using bowtie1^35^ in single-read mode using the default parameters. Transcripts without read-mapping support were removed. Functional annotation was performed using blastx v.2.2.31^36^ with an e-value < 1^-10^. NCBI RefSeq^37^, The Arabidopsis Information Resource (TAIR)^38^ and NCBI non-redundant (nr) databases were used as reference databases. We excluded transcripts that were exclusively similar to non-plant sequences. Protein family domains were identified using HMMER3^39^, which iteratively searched transcripts against the Pfam database. To assign Gene Ontology (GO) terms to transcripts, we used the *Arabidopsis thaliana* gene association file from the GO Consortium^40^ and retrieved the information for transcripts showing similar coding sequences in the *A. thaliana* genome. Redundant transcripts were clustered with Cd-hit-est v.4.6.1^41^ using local alignment mode with 95% identity and 70% coverage of the shortest sequence thresholds. Open reading frames (ORFs) were identified using Transdecoder (http://transdecoder.sf.net). We reduced the redundancy of transcripts in the final assembly by retaining for each CD-HIT-EST cluster either the sequence with the longest ORF or, in the absence of sequences containing ORF, the longest sequence.

The completeness of the final transcriptome was assessed using BUSCO^42^. Additionally, a reciprocal blastn alignment using an e-value threshold of 10^−10^ and a minimum alignment length of 100 nucleotides with at least 70% identity was used to compare the *A. schaueriana* transcriptome with other publicly available transcriptomes of congeneric species.

### Comparative transcriptomics using RNA-sequencing

Tissue-specific count data were obtained from the number of reads uniquely mapped to each transcript of the non-redundant transcriptome using bowtie1^35^ and normalised using edgeR^43^. Differentially expressed transcripts (DETs) between equatorial and subtropical tree tissue-specific samples were detected using the exact test for negative binomial distributions with an FDR < 0.05 (Benjamini-Hochberg). GO term enrichment analyses of the DETs were performed using GOseq^44^ with the Wallenius approximation method and P-value < 0.05. Differential expression results were verified using reverse transcription real-time PCR (qRT-PCR) (Supplementary Methods).

### Detection of candidate adaptive loci

We sampled leaves from 79 adult plants at ten locations spanning most of the distribution of *A. schaueriana* Stapf & Leechman ex Moldenke (Fig. 1, Supplementary information Table S2). We isolated DNA using the DNeasy Plant Mini Kit (QIAGEN) and NucleoSpin Plant II (Macherey Nagel) following the manufacturers’ instructions. DNA quality and quantity were assessed using 1% agarose electrophoresis and the QuantiFluor dsDNA System with a Quantus fluorometer (Promega). Nextera-tagmented reductively-amplified DNA (nextRAD) libraries^45^ were prepared and sequenced by SNPsaurus (SNPsaurus) on a HiSeq 2500 (Illumina, Inc.) with 100-bp single-end chemistry. Genomic DNA fragmentation and short adapter ligation were performed with the Nextera reagent (Illumina, Inc.) followed by amplification with one of the primers matching the adapter and extending nine arbitrary nucleotides into the genomic DNA. Assembly, mapping and single nucleotide polymorphic loci (SNP) identification were performed using custom scripts (SNPsaurus), which created a reference catalogue of abundant reads across the combined sample set and mapped reads to this reference, allowing two mismatches and retaining biallelic loci present in at least 10% of the samples. We further filtered markers by allowing no more than 65% of missing data, Phred score > 30, 8x minimum coverage, only one SNP per locus and a minor allele frequency ≥ 0.05 using vcftools v.0.1.12b^46^. To reduce paralogy or low-quality genotype calls, we used a maximum read coverage of 56 (the average read depth times 1.5 standard deviation).

After excluding plants morphologically identified as *A. schaueriana* with genomic signs of hybridisation with *A. germinans* (L.) L., we assessed the genetic structure considering all SNPs, using the discriminant analysis of principal components (DAPC)^47^ and ADMIXTURE v.1.3.0^48^. For DAPC analyses, we considered the number of groups (K) varying from 1-50 and the Bayesian information criteria for inferring K. Additionally, we used the optim.a.score function to avoid overfitting during the discrimination steps. For the ADMIXTURE analyses, we performed three separate runs for K varying from 1-15 using the block-relaxation method for point estimation; computing was terminated when estimates increased by < 0.0001, and the most likely K was determined by cross-validation.

We used two programs to minimise false-positive signs of natural selection: LOSITAN^49^, assuming an infinite allele model of mutation, using a confidence interval of 0.99, a false-discovery rate (FDR) of 0.1, the neutral mean FST and forcing the mean FST options; and pcadapt 3.0.4^50^, which simultaneously identifies the population structure and the loci excessively related to this structure, using an FDR < 0.1.

Putative evidence of selection was considered only for SNP loci that were conservatively identified by pcadapt and five independent runs of LOSITAN to avoid false-positives^51^. As selection is presumably stronger in coding regions of the genome and there is no reference genome for the species, we used the *de novo* assembled transcriptome, characterized herein, as a reference to identify candidate loci within putative coding regions. We performed a reciprocal alignment between nextRAD sequences (75 bp) and longer expressed sequences (≈ 300-11600 bp) using blastn v.2.2.31^36^, with a threshold of at least 50 aligned nucleotides, a maximum of one mismatch and no gaps.

### Data accessibility

Expression data and sequences that support the findings have been deposited in GenBank with the primary accession code GSE116060. A Variant Call Format file and its complementary file, both required for all of the genome-wide SNP diversity analyses are provided as Supplementary Files (Supplementary Datasets 1-2).

## Results

### Comparative physiology using a common garden experiment

Seedlings from equatorial and subtropical sites diverged in key ecophysiological traits in the common garden experiment (Fig. 2-4). The inclination angle of the leaf lamina and the average size of individual leaves were smaller in equatorial than in subtropical plants, but total leaf area and specific leaf area did not differ between the groups (Fig. 2, Supplementary information Table S3, Supplementary information Fig. S3). Additionally, equatorial plants showed lower stomatal conductance and transpiration rates than did subtropical plants (Fig. 4). Subtropical plants accumulated more biomass in leaves and roots than did equatorial plants. However, the stem dry mass ratio (MR) (stem dry biomass/plant dry biomass) was greater in equatorial plants, whereas the leaf MR (leaf dry biomass/plant dry biomass) was greater in subtropical plants (Fig. 2). The dry biomass accumulated in the stems and the root MR (root dry biomass/plant dry biomass) were indistinguishable between the groups (Supplementary information Table S3). Unexpectedly, 63% of the equatorial plants flowered after six months of growth (Supplementary information Fig. S3g). Since this was not observed in any subtropical plant, flowering plants were not used in the biomass allocation analyses.

**Figure 2.**
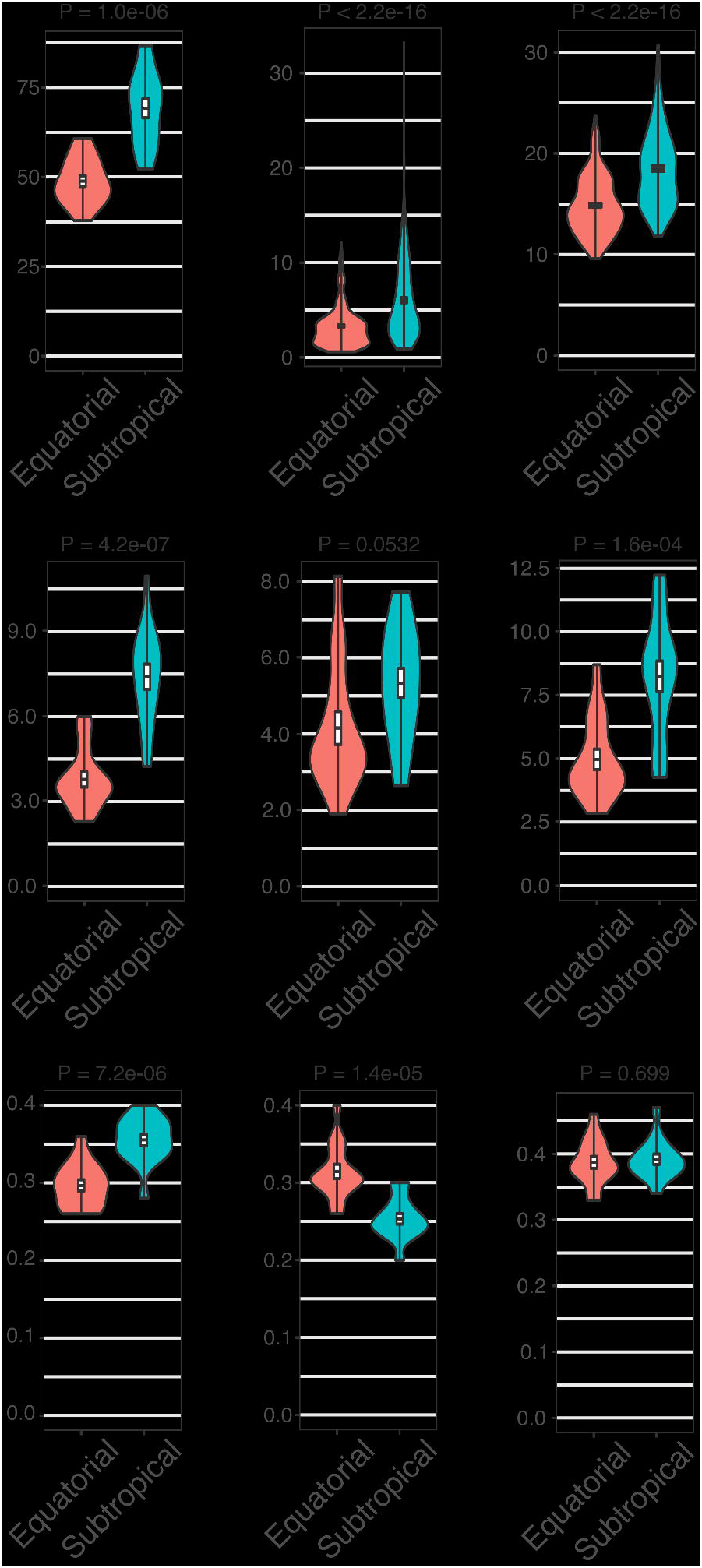
Pairwise comparison of morphological traits between *Avicennia schaueriana* seedlings from equatorial and subtropical sites grown using a common garden experiment. Violin plots represent the distribution of observations for plants from equatorial (red) and subtropical (blue) sampling sites. Box plots represent the mean, standard error, and maximum and minimum values. (a) Leaf inclination angle (n = 15 leaves per group, 5 plants per group); (b) individual leaf area (n = 250 leaves per group, 3 plants per group); (c) vessel lumen diameter (n = 180 vessels per group, 3 plants per group); (d-f) leaf, stem and root dry weight (n = 15 plants per group); and (g-i) leaf, stem and root dry mass ratio (organ-specific dry biomass/plant dry biomass) (n = 15 plants per group). For all variables represented in the figure, the absence of difference between groups was rejected using the unpaired Student’s T-test or the Mann-Whitney-Wilcoxon U-tests (b and c), at a significance threshold of 0.05.

**Figure 3.**
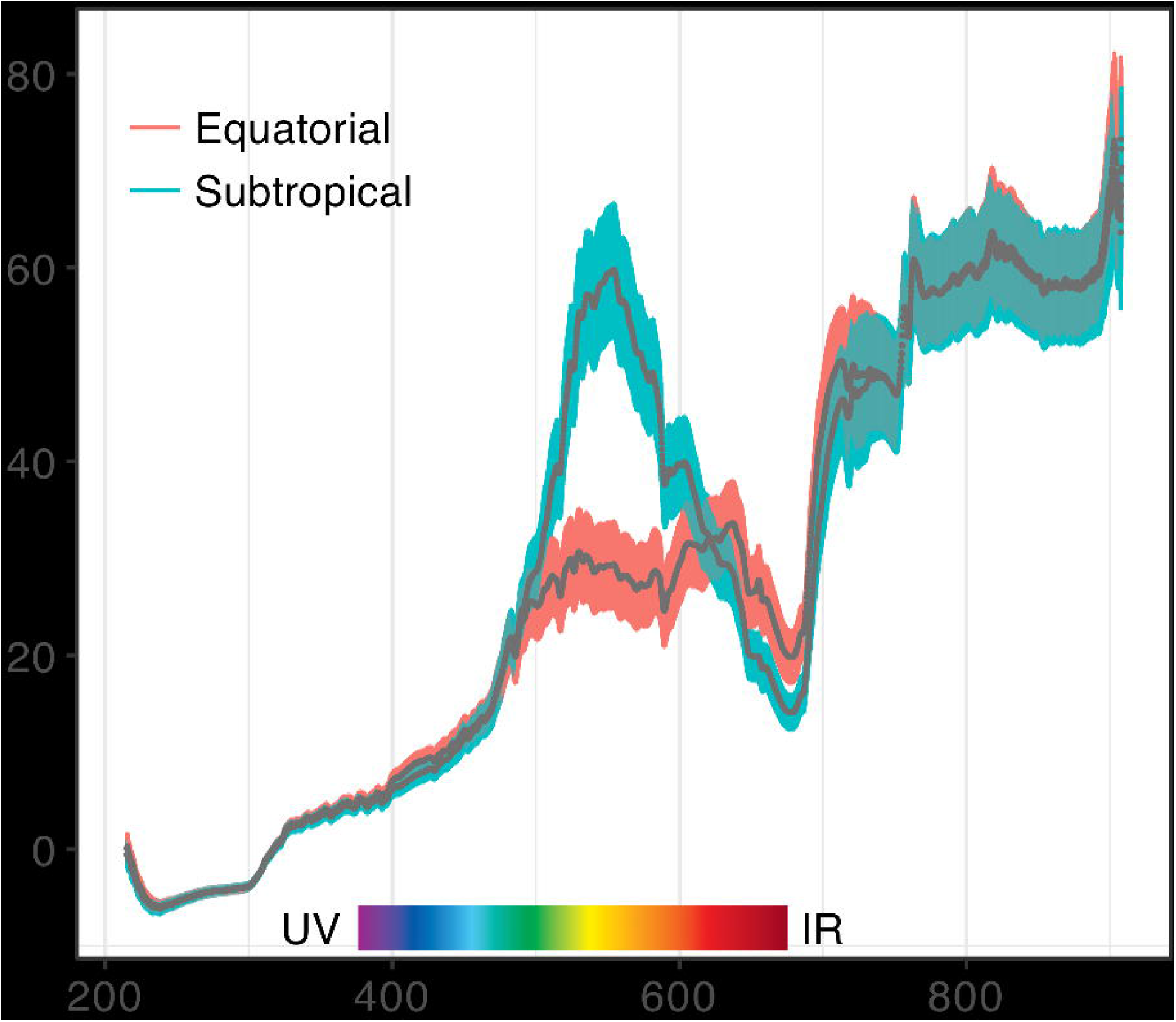
Light reflectance of the stem epidermis of five-month-old *Avicennia schaueriana* seedlings grown using a common garden experiment. Grey lines represent the mean reflectance, and colour-shaded areas represent the standard error for each seedling source site (red: equatorial; blue: subtropical, n = 10 plants per group). The visible light spectrum range is highlighted in the figure.

**Figure 4.**
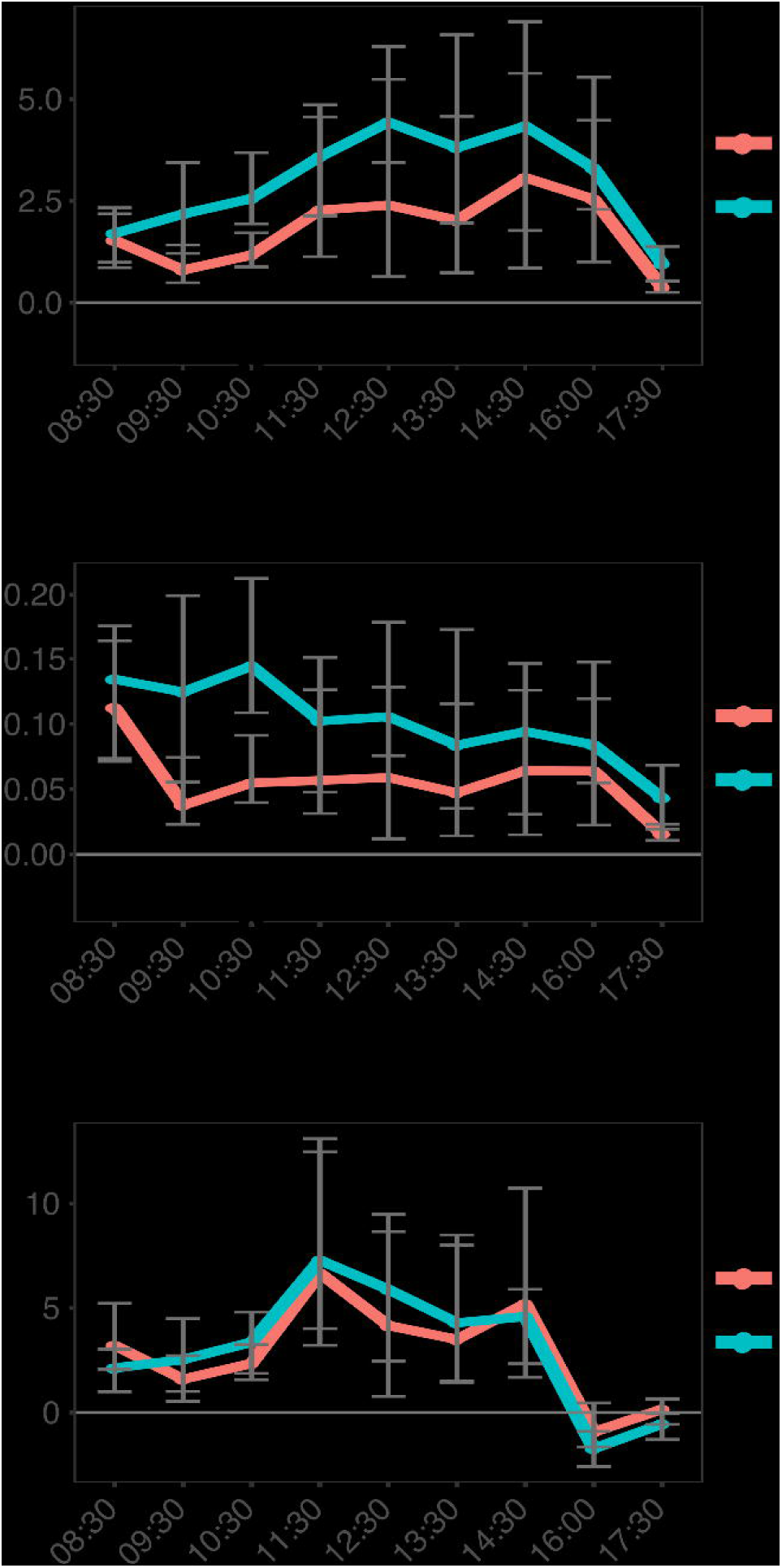
Daily curves of gas exchange in leaves from seven-months-old *Avicennia schaueriana* seedlings grown using a common garden experiment. * Represents rejection of the null hypothesis of the absence of a difference based on the unpaired Student’s t-test, at a significance threshold of 0.05 after Bonferroni correction (n = 10 plants per group). Error bars represent the standard error. Red line: mean values for equatorial samples; blue line: mean values for subtropical samples. (a) Transpiration rate; (b) stomatal conductance; (c) net CO_2_ assimilation rate.

Mean stem vessel diameter was smaller in the equatorial seedlings than in the subtropical seedlings, indicating enhanced hydraulic safety (Fig. 2, Supplementary information Fig. S3h-i). However, the vessel density, vessel grouping index, vessel lumen area in sapwood and total hydraulic conductivity of the stems were not significantly different between the groups (P-value > 0.05) (Supplementary information Table S3). Plants from contrasting latitudes exhibited different stem epidermis pigmentation, with equatorial seedlings reflecting more long wavelengths red light (635-700 nm) and less medium wavelengths green light (520-560 nm) than subtropical seedlings (Fig. 3).

### Characterisation of the <u>A. schaueriana</u> transcriptome

In the absence of a reference genome, we obtained the first reference transcriptome for *A. schaueriana* from leaves, stems and flowers of adult individuals under field conditions (Supplementary information Fig. S4, Supplementary information Table S1). Over 209 million paired-end 72-bp reads showing high quality were *de novo* assembled into a reference, non-redundant transcriptome containing 49,490 sequences, of which 30,227 (61%) were putative protein-coding sequences. Over 91.9% of these reads were mapped to a single transcript, indicating minimum redundancy and a wide representation of sequenced data (Supplementary information Table S4). Moreover, 91.8% of universal plant orthologous genes were present in the reference transcriptome (Supplementary information Table S4). Sequences with complete ORFs represented approximately 42% (12,796) of all putative protein-coding transcripts (Supplementary information Table S5, Supplementary information Fig. S5). Most of these putative protein-coding sequences (94.33%) showed significant similarity (e-value < 1e-10) to proteins in the Plant RefSeq and TAIR databases (Supplementary information Fig. S5c). More than 80% of protein-coding sequences matched proteins from *Sesamum indicum* or *Erythranthe guttata*, which, as *A. schaueriana*, belong to the order Lamiales (Supplementary information Fig. S6d). We identified 27,658, 18,325 and 13,273 putative orthologs between the *A. schaueriana* reference transcriptome and transcriptomes derived from *A. marina* leaves^52^ and *A. officinalis* leaves^53^ and roots^54^, respectively (Supplementary information Table S6).

### Comparative transcriptomics between trees from contrasting latitudes

To identify the effects of environmental variation on gene expression and, thus on phenotypes, at contrasting source sites in the field, we compared expression levels in trees under field conditions as comparable as possible. Sampling was conducted during the low tide, from 10:30 AM to 4:00 PM, at the end of winter at the subtropical site and at the beginning of the dry season at the equatorial site (Supplementary information Table S1). As expected, we observed a consistent source-site pattern in overall transcript expression levels from leaves and stems (Supplementary information Fig. S6a-b). However, flowers transcript expression levels did not show a clear pattern among samples of the same origin (Supplementary information Fig. S6c), leading to the identification of only one DET. Thus, we did not include flowers in the subsequent analyses (Supplementary information Fig. S6f). Conversely, 1,837 and 904 transcripts showed significantly different (FDR < 0.05) relative abundances between equatorial and subtropical samples in leaves and stems, respectively (Supplementary information Fig. S6d-e). Among the total 2,741 DETs, 1,150 (41.91%) were putative protein-coding transcripts.

The assignment of transcripts to GO terms was possible for 25,184 (83.31%) of the 30,227 putative protein-coding sequences, allowing GO enrichment analyses. Analyses were conducted separately for leaves and stems and for each of the following two sets of DETs: one showing higher expression levels in equatorial than in subtropical samples (which we refer to as “DET-Eq”) and the other showing higher abundance in subtropical than in equatorial samples (which are referred as “DET-St”). We focused on the biological processes associated with key aspects of the responses to contrasting climate conditions between the equatorial and subtropical sites (Table 1, Supplementary information Fig. S1). The enriched processes among the DET sets included photosynthesis; cell wall biosynthesis; plant responses to ultraviolet radiation (UV), temperature stimulus and water stress (Supplementary information Tables S7-S11, Supplementary information Fig. S6i-l).

#### Photosynthesis

Among the DET-St set, we observed various putative genes participating in the biosynthesis of the photosynthetic apparatus, chlorophyll and photoreceptors; the function of electron transporters and chloroplast movement coordination. Contrastingly, the DET-Eq set showed enrichment of transcripts similar to proteins required for disassembling the light-harvesting complex of photosystem II in thylakoid membranes^55^ (Supplementary information Table S11).

#### Cell wall biosynthesis

Transcripts similar to 33 distinct proteins and transcription factors that play central roles in the biosynthesis and deposition of cell wall components, such as cellulose, hemicellulose, lignin and pectin, were identified among DET-Eq (Supplementary information Table S10).

#### Response to UV

Both the DET-St and DET-Eq sets showed enrichment of functions related to responses to UV radiation; however, the transcript annotations differed between these sets. The DET-St set included putative UV-B protectants and genes involved in UV-B-induced antioxidant biosynthesis, such as plastid copper/zinc superoxide dismutases, photosystem II repair proteins, and L-ascorbic acid. In contrast, the DET-Eq set showed enrichment of transcripts associated with photo-oxidative damage reduction and the positive regulation of anthocyanin biosynthesis in response to UV. Antioxidants induced by UV irradiation^56^, such as putative iron superoxide dismutases and pyridoxine biosynthesis genes, were among the DET-Eq set (Supplementary information Table S11).

#### Response to temperature

In the DET-St set, we observed putative genes presenting critical roles in chloroplast protein translation during cold acclimation and that provide tolerance to chilling temperatures^57, 58^. There included transcripts similar to the *GLYCINE-RICH RNA-BINDING* (RZ1A), which has a chaperone activity during cold acclimation^59^, and to the cold-inducible ATP-dependent *DNA HELICASE* ATSGS1, required for DNA damage-repair^60^. Interestingly, DET-St included a putative *AGAMOUS-LIKE 24* (AGL24) transcription factor that is involved in vernalisation-induced floral transition^61^. Contrastingly, various transcripts similar to heat shock-inducible chaperones and to *ADENINE NUCLEOTIDE ALPHA HYDROLASE-LIKE* (AT3G53990), involved in chaperone-mediated protein folding^62^, were among the DET-Eq set, potentially enhancing tolerance to heat in equatorial plants. Additionally, a transcript similar to the *ETHYLENE-RESPONSIVE ELEMENT BINDING* (RAP2.3), which confers resistance to heat and hydrogen peroxide^63^, was observed in this group (Supplementary information Table S11).

#### Response to water stress

Transcripts associated with the response and tolerance to water deficits and with cellular ion homeostasis and osmotic adjustment were enriched among DET-Eq; for instance, a transcript similar to the *ETHYLENE-RESPONSIVE TRANSCRIPTION FACTOR* (RAP2.4), which regulates the expression of several drought-responsive genes, including aquaporins^64, 65^. Accordingly, a putative aquaporin *PLASMA MEMBRANE INTRINSIC* (PIP1;4)^66^ was found in this set. In DET-Eq, we observed putative genes participating in the synthesis of raffinose, an osmoprotective soluble trisaccharide^67^, as well as transcripts similar to osmosensitive ion channels belonging to the *EARLY-RESPONSIVE TO DEHYDRATION STRESS FAMILY*. Correspondingly, we observed an ion channel, *SLAC1 HOMOLOGUE 3* (SLAH3), required for stomatal closure induced by drought stress^68^, and the putative *NINE-CIS-EPOXYCAROTENOID DIOXYGENASE 3* (NCED3), which increases plant resistance to water deficits through the accumulation of abscisic acid (ABA), leading to stomatal closure. Possibly as a consequence of decreased stomatal conductance, a transcript similar to the photorespiration gene, *ALANINE-GLYOXYLATE AMINOTRANSFERASE 2 HOMOLOG 3* (AT3G08860)^69^, was also observed among the DET-Eq (Supplementary information Table S11).

We confirmed the results obtained from RNA-Seq data computational analyses by qRT-PCR using ten DETs from leaf samples (Supplementary information Fig. S7, Supplementary information Table S12, Supplementary Note).

### Detection of SNPs with signs of selection

RNA-Seq enabled the assembly of a reference transcriptome and the identification of biological processes influenced by the environmental divergence over the South American coastline. To complement these analyses, as one cannot disentangle the effects of plasticity and adaptive selection in the differential expression^70, 71^, we searched for gene sequence variations among trees sampled along the Atlantic coast of South America (Fig. 1, Supplementary information Table S2). After quality filtering of the sequenced data, we selected 77 individuals without evidence of interspecific hybridisation with *A. germinans* for downstream analyses. We identified a set of 6,170 high-quality unlinked biallelic SNP loci with a minor allele frequency ≥ 0.05 and ≥ 8x coverage. The overall genetic structure of genome-wide SNPs corroborated a previous study based on putatively neutral loci^22^, dividing the species into two populations: north and south of the northeast extremity of South America (Fig. 5).

**Figure 5.**
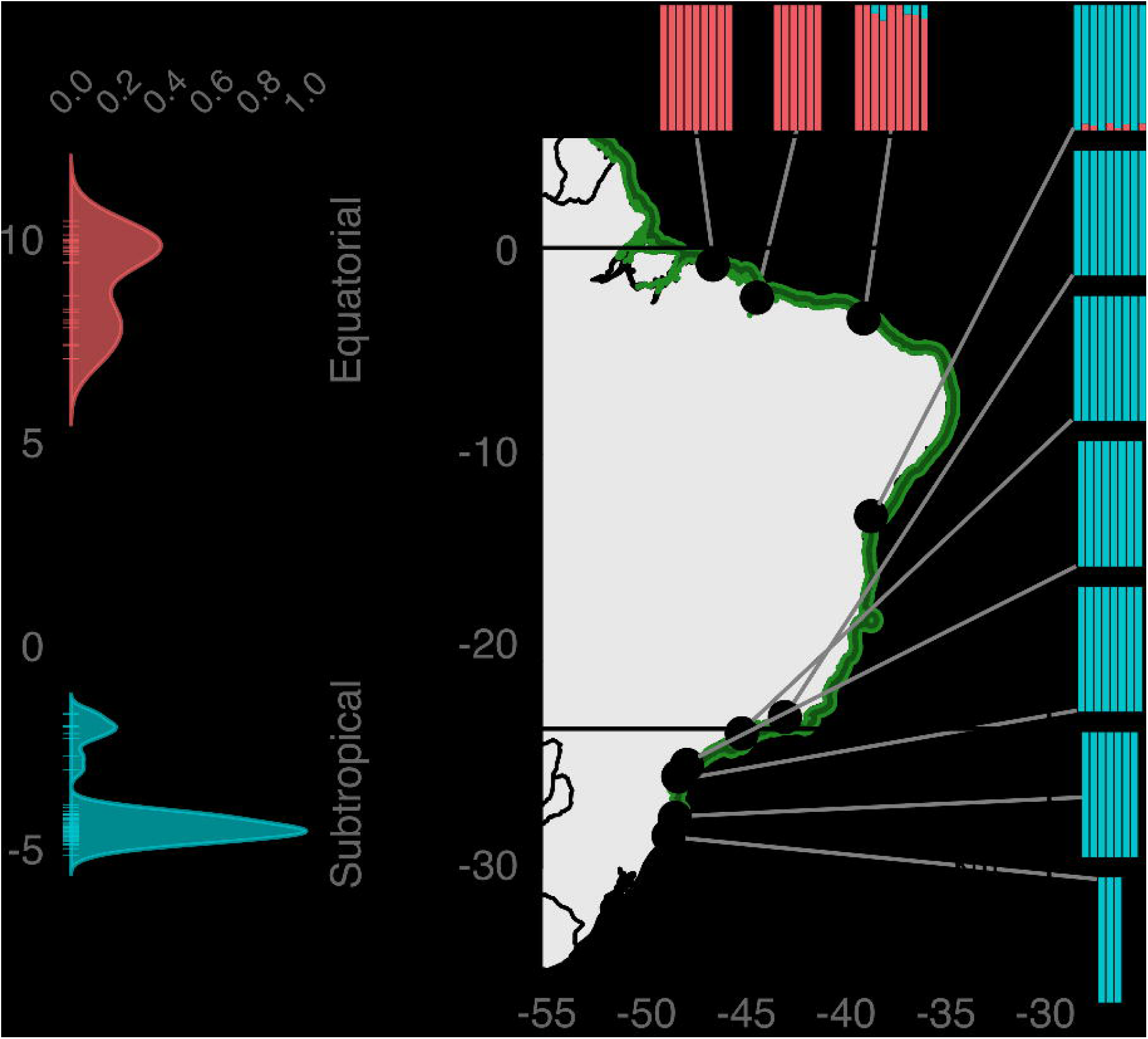
Population structure of *A. schaueriana* inferred from genome-wide genotyping of single nucleotide polymorphic (SNP) loci. (a) Density of individuals on the first retained discriminant function, calculated using a multivariate method, the discriminant analysis of principal components (DAPC); different colours represent inferred populations; (b) map showing the distribution of the species (green shaded area), the geographical location of sampling sites (points) and barplots in which each colour denotes an ancestral cluster and each bar represents an individual as inferred by the model-based method implemented in Admixture 1.3.0.

We observed 122 loci showing considerable departures from neutral expectations of interpopulational divergence, as conservatively detected^72^ by pcadapt and LOSITAN. Fifteen of these loci aligned to *A. schaueriana* transcripts that were similar to gene models in *A. thaliana* and *S. indicum* (Supplementary information Table S13), enabling screening for their potential functional roles. However, five reference proteins did not have informative functions described for the model plant, hindering functional inferences. Conversely, among the remaining annotated candidates, we found five putative genes involved in processes related to contrasting equatorial and subtropical environmental variables (Fig. 6). One candidate locus was detected in the putative transcription factor *RAP2.4*, which is induced in response to water and salt stress^64^ and regulates the expression of aquaporins^65^. Two other candidates showed similarity to the transcription factors *ZINC-FINGER PROTEIN 1* (ZFN1), involved in the regulation of the expression of several water stress genes^73^, and the *HYPOXIA RESPONSE ATTENUATOR* (HRA1), which is strongly induced in response to low oxygen levels^74^. A putative *UDP GLUCOSYL TRANSFERASE*, an enzyme that catalyses the final step of anthocyanin biosynthesis wherein pigmentation is produced^75^, also showed evidences of positive selection. Additionally, one candidate locus was in a transcript similar to the *TETRATRICOPEPTIDE REPEAT* gene (AT2G20710, TPR), which might play a role in the biogenesis of the photosynthetic apparatus^76^.

**Figure 6.**
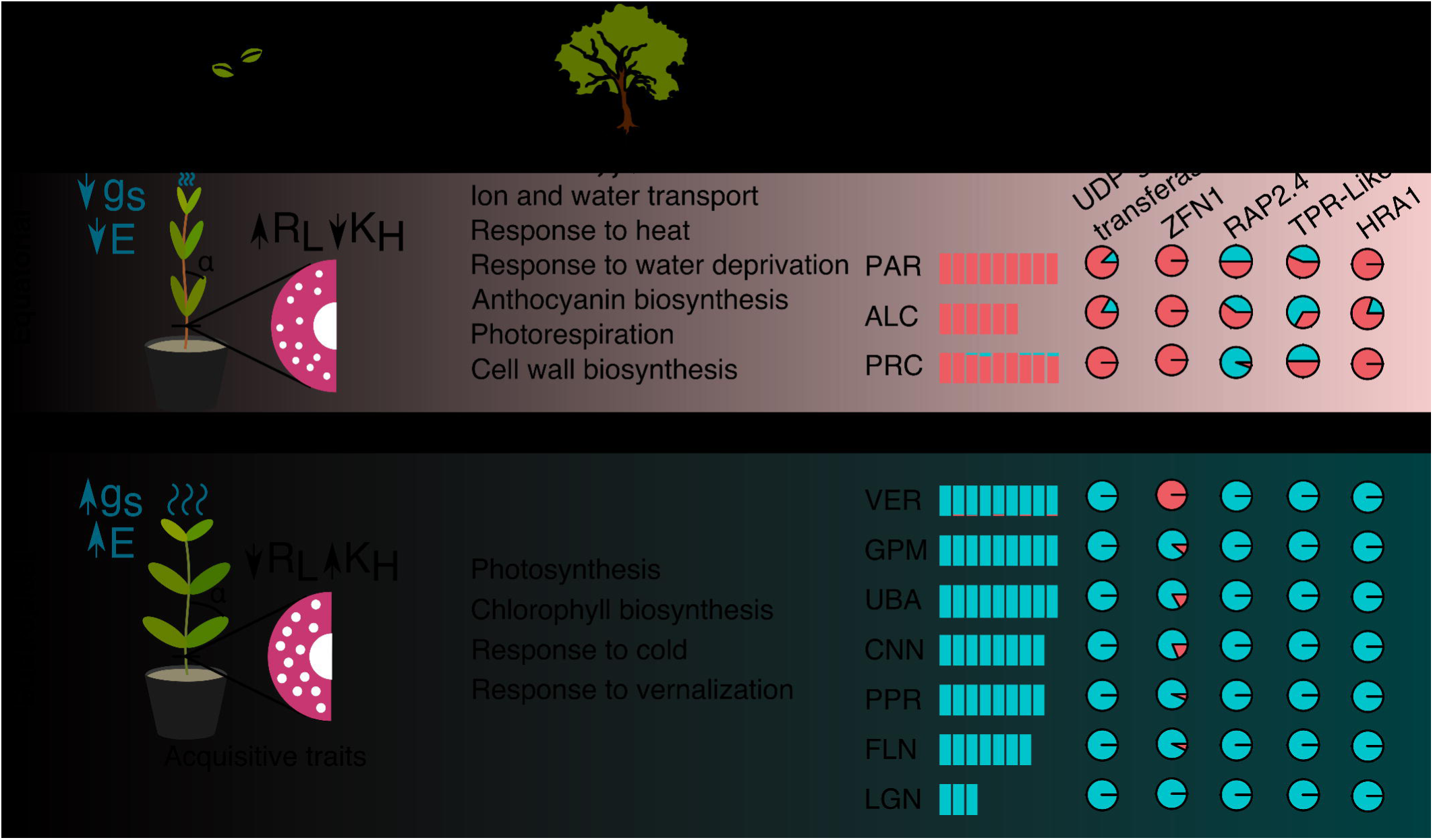
Graphical summary of results from ecophysiological and molecular approaches performed in this work. Under homogeneous conditions, plants from contrasting latitudes showed key divergences related to the use of water and to the acquisition of carbohydrates in ecophysiological traits. Differences associated with the response to contrasting environmental conditions were detected in transcripts expression levels of trees sampled in their natural habitats. Additionally, a north-south genetic divergence was observed in 77 trees sampled along the latitudinal gradient using genotypic data from over 6 thousand polymorphic loci. We also identified signatures of differential selective pressures on specific loci associated with the accumulation of anthocyanins (UDP-GLUCOSYLTRANSFERASE), with photosynthesis (TPR), and with the response to osmotic stress (RAP2.4 and ZFN1) and to hypoxia (HRA1). Our results suggest that the establishing success of propagules over the latitudinal gradient of the Atlantic coast of South America may play a role in shaping the diversity of genotypes and phenotypes in the widespread mangrove, *Avicennia schaueriana*. E: transpiration rate; g_S_: stomatal conductance; R_L_: xylem vessel lumen resistivity; K_H_: xylem vessel conductivity.

## Discussion

In this study, we integrated genomic and ecophysiological approaches to investigate the foundations of adaptive variations in a dominant tree from the Atlantic coast of South America. We tested the hypothesis that latitudinal variations in climate regimens shape genetic and phenotypic variation involved in resource-use strategies in widespread coastal trees, using the mangrove *Avicennia schaueriana* as a model. Overall, our results supported this hypothesis. Using a common garden experiment, we detected differences between plants from contrasting latitudes in key ecophysiological traits involved in carbon and water balances, indicating an inheritable basis of trait divergence. Accordingly, transcriptomic changes between plants sampled in contrasting latitudes showed enrichment in processes associated with central aspects of the environmental divergence across latitudes, such as responses to temperature, solar radiation, water deficit, photosynthesis and cell wall biosynthesis (Supplementary information Fig. S6i-l). The relevance of these biological processes in the field was corroborated by the identification of SNP loci with signs of selection present within putative coding transcripts similar to genes involved in responses to osmotic stress, accumulation of anthocyanins and photosynthesis, unveiling a genetic basis for environmental adaptation. Figure 6 summarises the integration of three independent approaches that converge to suggest population adaptation to contrasting environments over the species range.

### Evidence of a conservative resource-use strategy in the equatorial population of <u>A. schaueriana</u>

Traits exhibited by equatorial plants compared to subtropical plants during the common garden experiment, such as a smaller leaf size and angle, higher levels of red light-reflecting pigments, narrower vessels and lower rates of stomatal conductance, limit carbon acquisition^77^ and may have imposed constraints on carbon gain in equatorial plants. Accordingly, these plants also accumulated less biomass than subtropical plants (Fig. 2-4, Supplementary information Fig. S3). Despite causing limitations in growth, these characteristics allow plants to maintain suitable leaf temperature for photosynthesis while reducing the risk of cavitation, UV exposure and water loss through the minimisation of evaporative cooling^77^. Given the relevance of these traits to water and carbon balances, especially under high salinity environments, the prevalence of more conservative resource-use traits among equatorial samples in the common garden experiment suggested a selection favouring drought tolerance, likely being advantageous during hot-dry seasons in equatorial wetlands (August-December). These seasons frequently present a combination of stressful conditions, including high UV exposure; highly fluctuating soil salinity, resulting from wide daily tidal range; and high vapour pressure deficit (VPD), resulting from high temperature (> 30 °C) and an air humidity below 60% (Figure 1, Table 1, Supplementary information Fig. S1). Accordingly, equatorial plants also showed lower transpiration rates than subtropical plants in the common garden (Fig. 4). Additionally, 63% of the six-month-old equatorial plants flowered from July-August in the common garden experiment (Supplementary information Fig. S3g), whereas subtropical plants did not flower. This period is consistent with phenological observations reported for *A. schaueriana* in both equatorial^78^ and southern subtropical forests^79^, and could indicate a genetic basis for the observed variation. Early flowering is a phenotype with complex genetic components, rarely studied in non-model organisms, but is renowned as an adaptive mechanism for maximising chances of reproduction under stressful environments^80^.

In their native environment, equatorial plants showed increased expression levels of putative heat-shock proteins, drought-induced ion transporters, aquaporins and genes that play central roles in stomatal closure and photorespiration compared to subtropical plants, likely contributing to heat and drought tolerance. These findings provided multiple lines of evidence of heat and water stress responses. Higher expression of aquaporins and genes involved in the accumulation of organic solutes may contribute to lowering the cellular water potential, while improving drought tolerance in equatorial plants compared to subtropical plants^67, 81^. Additionally, higher expression of several transcripts associated with secondary cell wall biosynthesis and thickening, may enhance the rigidity of cells and reduce the risk of collapse during dehydration-rehydration cycles in equatorial trees^82^. Moreover, equatorial plants showed lower expression of several putative photosynthesis genes than subtropical plants. In response to drought, high-light and heat stress plants minimise the photooxidative damage by reducing the photosynthetic activity via repression of photosynthesis genes^83–86^. Remarkably, evidences of selection were detected in two putative transcription factors, *RAP2.4* and *ZFN1*, which play key roles in the regulation of osmotic stress-response^64, 73^. These findings support the hypothesis that dry seasons marked by low rainfall and high VPD, which are caused by the combination of high temperatures and low air humidity^87^ in the equatorial region, induce a pivotal selective pressure for coastal trees populations.

### Evidence of an acquisitive resource-use strategy in the subtropical population of <u>A. schaueriana</u>

Subtropical plants showed higher stomatal conductance and transpiration rates, higher levels of green light-reflecting pigments, larger leaf area, wider leaf lamina angle and larger xylem vessel diameter than equatorial plants in the common garden experiment (Fig. 2-4, Supplementary information Fig. S3). These characteristics enhance light energy absorbance and carbon acquisition, at the expense of a higher cavitation risk^88, 89^. Conversely, this apparent riskier strategy may compensate for seasonal declines in growth resulting from decreasing temperature, photoperiod and light quality at higher-latitudes^90^ (Table 1, Supplementary information Fig. S1). Although low temperatures reduce enzymatic activity^91^ and, thus, plant growth, the severity of low-temperature stress in the southernmost subtropical mangrove forests of the Americas is likely insufficient to favour the selection of freezing-tolerant adaptations, in contrast to results reported for mangroves at their northernmost edge on the Atlantic coast of the Americas^92^. At the southernmost limit of American mangrove forests, the minimum air temperature does not drop below 0 °C (Table 1, Supplementary information Fig. S1) and remains within the expected mangrove physiological thresholds^93^. Additionally, the small population size of *A. schaueriana* at this location^27^ and the arrival of maladapted propagules from northerly populations likely reduce the potential strength of selection favouring cold-adaptive traits.

Corroborating the observed differences in morphophysiological traits, we found divergence at the molecular level that may also be related to the increasing amplitude of variation in photoperiod and light quality towards high latitudes. Plants optimise the use of light energy and adjust photosynthetic activity through the regulation of light-harvesting and photosystem-component genes^83^. Thus, the higher expression levels of transcripts associated with photosynthesis in subtropical than in equatorial plants may have facilitated the absorption of light energy in subtropical plants during winter. Although solar irradiance levels were indistinguishable between sampling sites at the time of sampling (Supplementary information Table S1), transcriptomic changes in putative UV-inducible antioxidant and photodamage repair genes suggest the use of distinct strategies to respond to differential seasonality in photoperiod and solar radiation levels between subtropical and equatorial latitudes. Notably, we observed signatures of selection in two transcripts, one showing similarity to a *UDP-GLUCOSYL TRANSFERASE*, a key enzyme in the anthocyanin biosynthesis pathway, and the other in a transcript similar to a TPR gene, required for chlorophyll biosynthesis and for the biogenesis of the photosynthetic apparatus^76^. These results imply that solar radiation, in addition to VPD, may act as an environmental factor driving selection in *A. schaueriana*.

Although soil oxygen availability affects plant growth in intertidal areas, we did not focus our experiments on the relevance of hypoxia in shaping adaptive divergence in coastal wetlands^94^. However, evidence of selection was detected in a transcription factor highly induced by oxygen deprivation (HRA1)^74^. Additionally, the HRA1 putative homolog also showed a 1.75-fold higher expression in subtropical leaves relative to that in equatorial leaves (Supplementary information Table S1), even though sampling was conducted during the low tide at both sites. As tidal amplitude decreases with increasing latitude along the Atlantic coast of South America^95^ (Table 1, Fig. 6), trees are exposed to increasing anoxia conditions southwards. These findings suggest that subtropical populations may have better stress preparedness for hypoxia than equatorial populations.

### Climate change and conservation implications

Our results provide compelling evidence that adaptations to contrasting seasonality in freshwater availability and solar radiation are involved in the organisation of diversity of the dominant coastal tree *A. schaueriana*. The functional divergence described herein might differentially affect the sensitivity of northerly and southerly populations to a rapidly changing climate. It has been suggested that the southernmost mangrove forests in the Americas could expand polewards in the near future^96^. We expect that the observed acquisitive resource-use characteristics may indeed favour subtropical tree growth under increased CO_2_ concentrations, temperatures and rainfall, as predicted for this region by 2100^97^. However, a great landward relocation of subtropical populations will be necessary to their persistence in the face of rising sea level^98^, due to the narrowing of intertidal zones towards mid-latitude areas on the Atlantic coast of South America (Fig. 1). Even though subtropical populations apparently show a better preparedness for hypoxia than equatorial populations, this scenario is aggravated by the dense coastal occupation by urban and agricultural areas, which may preclude the landward migration of subtropical mangroves in South America. Contrastingly, equatorial populations frequently have wider plains that are potentially available for expansion and lower human occupation, but might be threatened by reduced rainfall and increased VPD during more intense El Niño-Southern Oscillation events^97^. Increased temperature will stimulate both respiration^99^ and photorespiration^100^ and may eventually offset benefits in carbon acquisition caused by increased CO_2_ concentrations^101^. The critical temperature threshold for photosynthesis may be overcome more frequently, possibly reducing carbon assimilation and productivity^102^. With a conservative resource-use strategy, further limitations in net carbon assimilation could, in extreme events, lead to biomass loss or tree mortality triggered by carbon starvation^103^. This scenario could be especially severe to the semiarid northeast South American coast, particularly in the face of intense human use of adjacent plains and decreasing trends in terrestrial freshwater availability and coastal freshwater input^104, 105^.

Our results corroborate previous studies of marine species from the Northern Atlantic that associated limited north-south population connectivity with adaptation to latitudinal environmental dissimilarity^19–21^. As widespread trees distributed along the Atlantic coast of South America show an overlapping north-south organisation of genetic variation^22–24^, we hypothesise that they may also show divergent adaptations to heterogeneous resource availability over their distribution ranges. Conservation efforts of coastal ecosystems should focus on resilient, habitat-forming species that give shelter and act as climate rescuers for several stress-sensitive species^106^. We recommend that populations of coastal trees occurring north and south from the northeast extremity of South America should warrant attention as distinct conservation management units^107^ for the long-term persistence of coastal biodiversity and the ecosystem services they provide.

## Conclusions

Studies on local adaptations in tropical trees are frequently limited by difficulties in implementing traditional approaches, such as reciprocal transplants, due to their long generation time and the lack of basic biological information, as knowledge of their evolutionary history, reproductive mode or phenological patterns. Despite limitations, we provided integrated ecophysiological and genomic evidences that support the hypothesis that seasonal environmental heterogeneity in solar radiation and freshwater availability may shape the variation of traits associated with resource-use in *A. schaueriana*, a widespread tree along the Atlantic coastline of South America. The non-neutral trait diversity derived from the interdisciplinary approach used here provides a perspective into the molecular and phenotypic scales at which environmental selection may shape functional variation of this dominant species, on a continental scale. Integrating high-throughput sequencing and ecophysiological approaches, has shown to be a promising strategy in the study of adaptation in non-model, long-lived species. To provide more realistic predictions of how dominant coastal trees may respond to current global climate changes we encourage the development of further studies accounting for phenotypically^14^ and genetically^108^ informed distribution modelling. Since freshwater availability has been decreasing in coastal areas worldwide^104^, strongly compromising the productivity of coastal plants^105^, such studies should focus on biological variation involved in the balance between carbon gain and water loss. This knowledge can improve predictions on the future of ecosystems they form and generate key information for forest conservation and management efforts^14, 109^. Without the realization and dissemination of such studies, the success of conservation plans for tropical forests and their potential to mitigate climate change will likely be seriously compromised.

## Acknowledgements

We thank Prof. Michel G. Vincentz and Prof. Juliana S. Mayer for advising and making their laboratory available for our training and performing of RNA extraction and slide preparation for the wood anatomy analyses. We acknowledge Ilmarina C. Menezes and Stephanie K. Bajay for assistance in the field. We thank Stephanie K. Bajay for providing assistance in the germination of the propagules used in the common garden experiment. Moreover, we thank Greg Mellers and Alessandro Alves-Pereira for critically reading the manuscript. M.V.C. received fellowships from the São Paulo Research Foundation - FAPESP (PhD 2013/26793-7) and from the Coordination for the Improvement of Higher Education Personnel - CAPES Computational Biology Program (PhD SWE 8084/2015-07, PhD 88887.177158/2018-00). G.M.M. received fellowships from FAPESP (PD 2013/08086-1, PD BEPE 2014/22821-9) and a grant from the Brazilian National Council for Scientific and Technological Development - CNPq (PD 448286/2014-9). C.C.S. received fellowships from FAPESP (PD 2015/24346-9) and CAPES Computational Biology Program (88881.185134/2018-01). R.S.O. received a CNPq productivity scholarship and a grant from Microsoft research-FAPESP (2011/52072-0). A.P.S. received a grant from CAPES - Computational Biology Program (88882.160095/2013-01), a research fellowship from CNPq (309661/2014-5) and a CNPq productivity scholarship.

## Author contributions

A.P.S., R.S.O., G.M.M. and M.V.C. designed the study. M.V.C. and G.M.M. conducted fieldwork. M.V.C. and C.S.M. cultivated seedlings and performed analyses of morphophysiological data. M.V.C. prepared samples and performed RNA-sequencing. M.V.C., M.D., D.H.O. and G.M.M. analysed RNA-Seq data. C.C.S. and M.V.C. verified RNA-Seq data through qRT-PCR. G.M.M. prepared samples and performed genotyping of genome-wide SNP. G.M.M. and M.V.C. analysed nextRAD results. A.P.S., M.I.Z., G.M.M. and R.S.O. contributed material/reagents/analysis tools. M.V.C. and G.M.M. wrote the manuscript. All authors discussed the results and contributed to the manuscript.

## References

1. Hereford, J. A Quantitative Survey of Local Adaptation and Fitness Trade-Offs. Am. Nat. 173, 579–588 (2009).

2. Kawecki, T. J. & Ebert, D. Conceptual issues in local adaptation. Ecol. Lett. 7, 1225–1241 (2004).

3. Albert, C. H., Grassein, F., Schurr, F. M., Vieilledent, G. & Violle, C. When and how should intraspecific variability be considered in trait-based plant ecology? Perspect. Plant Ecol. Evol. Syst. 13, 217–225 (2011).

4. Aitken, S. N., Yeaman, S., Holliday, J. A., Wang, T. & Curtis-McLane, S. Adaptation, migration or extirpation: climate change outcomes for tree populations. Evol. Appl. 1, 95–111 (2008).

5. Jump, A. S. & Peñuelas, J. Running to stand still□: adaptation and the response of plants to rapid climate change. Ecol. Lett. 8, 1010–1020 (2005).

6. Hoegh-Guldberg, O. & Bruno, J. F. The Impact of Climate Change on the World’s Marine Ecosystems. Science (80-.). 328, 1523–1528 (2010).

7. Loarie, S. R. et al. The velocity of climate change. Nature 462, 1052–1055 (2009).

8. Duke, N. C. et al. A World Without Mangroves? Science (80-.). 317, 41 (2007).

9. Polidoro, B. A. et al. The loss of species: Mangrove extinction risk and geographic areas of global concern. PLoS One 5, e10095 (2010).

10. Normile, D. El Niño’s warmth devastating reefs worldwide. Science (80-.). 352, 15–16 (2016).

11. Duke, N. et al. Large-scale dieback of mangroves in Australia. Mar. Freshw. Res. 68, 1816–1829 (2017).

12. Marbà, N. & Duarte, C. M. Mediterranean warming triggers seagrass (*Posidonia oceanica*) shoot mortality. Glob. Chang. Biol. 16, 2366–2375 (2010).

13. Kauffman, J. B. et al. Carbon stocks of mangroves and salt marshes of the Amazon region, Brazil. Biol. Lett. 14, 20180208 (2018).

14. Moran, E. V., Hartig, F. & Bell, D. M. Intraspecific trait variation across scales: Implications for understanding global change responses. Glob. Chang. Biol. 22, 137–150 (2016).

15. Wee, A. K. S. et al. The integration and application of genomic information in mangrove conservation. Conserv. Biol. 33, 206–209 (2018).

16. Takayama, K., Tamura, M., Tateishi, Y., Webb, E. L. & Kajita, T. Strong genetic structure over the American continents and transoceanic dispersal in the mangrove genus *Rhizophora* (Rhizophoraceae) revealed by broad-scale nuclear and chloroplast DNA analysis. Am. J. Bot. 100, 1191–1201 (2013).

17. Mori, G. M., Zucchi, M. I., Sampaio, I. & Souza, A. P. Species distribution and introgressive hybridization of two *Avicennia* species from the Western Hemisphere unveiled by phylogeographic patterns. BMC Evol. Biol. 15, 1–15 (2015).

18. Marshall, D. J., Monro, K., Bode, M., Keough, M. J. & Swearer, S. Phenotype-environment mismatches reduce connectivity in the sea. Ecol. Lett. 13, 128–140 (2010).

19. Jeffery, N. W. et al. Range-wide parallel climate-associated genomic clines in Atlantic salmon. R. Soc. Open Sci. 4, 171394 (2017).

20. Bradbury, I. R. et al. Parallel adaptive evolution of Atlantic cod on both sides of the Atlantic Ocean in response to temperature. Proc. R. Soc. B Biol. Sci. 277, 3725–3734 (2010).

21. Chu, N. D., Kaluziak, S. T., Trussell, G. C. & Vollmer, S. V. Phylogenomic analyses reveal latitudinal population structure and polymorphisms in heat stress genes in the North Atlantic snail *Nucella lapillus*. Mol. Ecol. 23, 1863–1873 (2014).

22. Mori, G. M., Zucchi, M. I. & Souza, A. P. Multiple-geographic-scale genetic structure of two mangrove tree species: the roles of mating system, hybridization, limited dispersal and extrinsic factors. PLoS One 10, e0118710 (2015).

23. Francisco, P. M., Mori, G. M., Alves, F. M., Tambarussi, E. & Souza, A. P. De. Population genetic structure, introgression, and hybridization in the genus *Rhizophora* along the Brazilian coast. Ecol. Evol. 8, 3491–3504 (2018).

24. Takayama, K., Tateishi, Y., Murata, J. & Kajita, T. Gene flow and population subdivision in a pantropical plant with sea-drifted seeds *Hibiscus tiliaceus* and its allied species: Evidence from microsatellite analyses. Mol. Ecol. 17, 2730–2742 (2008).

25. Savolainen, O., Pyhäjärvi, T. & Knürr, T. Gene Flow and Local Adaptation in Trees. Annu. Rev. Ecol. Evol. Syst. 38, 595–619 (2007).

26. Barrett, R. D. H. & Hoekstra, H. E. Molecular spandrels: tests of adaptation at the genetic level. Nat. Rev. Genet. 12, 767–780 (2011).

27. Soares, M. L. G., Estrada, G. C. D., Fernandez, V. & Tognella, M. M. P. Southern limit of the Western South Atlantic mangroves: Assessment of the potential effects of global warming from a biogeographical perspective. Estuar. Coast. Shelf Sci. 101, 44–53 (2012).

28. Souza-Filho, P. W. M. et al. Holocene coastal evolution and facies model of the Bragança macrotidal flat on the Amazon Mangrove Coast, northern Brazil. J. Coast. Res. 1, 306–310 (2006).

29. Kjerfve, B. et al. Morphodynamics of muddy environments along the Atlantic coasts of North and South America. in Muddy coasts of the World: Processes, Deposits and Function (eds. Healy, T., Wang, Y. & Healy, J.-A.) 4, 479–532 (Elsevier Science B.V., 2002).

30. Reef, R. et al. The effect of atmospheric carbon dioxide concentrations on the performance of the mangrove *Avicennia germinans* over a range of salinities. Physiol. Plant. 154, 358–368 (2015).

31. Schneider, C. A., Rasband, W. S. & Eliceiri, K. W. NIH Image to ImageJ: 25 years of image analysis. Nat. Methods 9, 671–675 (2012).

32. Scholz, A., Klepsch, M., Karimi, Z. & Jansen, S. How to quantify conduits in wood? Front. Plant Sci. 4, 1–11 (2013).

33. Oliveira, R. R., Viana, A. J. C., Reátegui, A. C. E. & Vincentz, M. G. A. An efficient method for simultaneous extraction of high-quality RNA and DNA from various plant tissues. Genet. Mol. Res. 14, 18828–18838 (2015).

34. Patel, R. K. & Jain, M. NGS QC toolkit: A toolkit for quality control of next generation sequencing data. PLoS One 7, e30619 (2012).

35. Langmead, B., Trapnell, C., Pop, M. & Salzberg, S. Ultrafast and memory-efficient alignment of short DNA sequences to the human genome. Genome Biol. 10, R25 (2009).

36. Camacho, C. et al. BLAST+: architecture and applications. BMC Bioinformatics 10, 421 (2009).

37. O’Leary, N. A. et al. Reference sequence (RefSeq) database at NCBI: Current status, taxonomic expansion, and functional annotation. Nucleic Acids Res. 44, D733–D745 (2016).

38. Berardini, T. Z. et al. The Arabidopsis Information Resource: Making and Mining the “Gold Standard” Annotated Reference Plant Genome. Genesis 53, 474–485 (2015).

39. Finn, R. D. et al. Pfam: The protein families database. Nucleic Acids Res. 42, 222–230 (2014).

40. Blake, J. A. et al. Gene ontology consortium: Going forward. Nucleic Acids Res. 43, D1049–D1056 (2015).

41. Li, W. & Godzik, A. Cd-hit: A fast program for clustering and comparing large sets of protein or nucleotide sequences. Bioinformatics 22, 1658–1659 (2006).

42. Simão, F. A., Waterhouse, R. M., Ioannidis, P., Kriventseva, E. V. & Zdobnov, E. M. BUSCO: Assessing genome assembly and annotation completeness with single-copy orthologs. Bioinformatics 31, 3210–3212 (2015).

43. Robinson, M. D., McCarthy, D. J. & Smyth, G. K. edgeR: a Bioconductor package for differential expression analysis of digital gene expression data. Bioinformatics 26, 139–140 (2010).

44. Young, M. D., Wakefield, M. J., Smyth, G. K. & Oshlack, A. Gene ontology analysis for RNA-seq: accounting for selection bias. Genome Biol. 11, R14 (2010).

45. Russello, M. A., Waterhouse, M. D., Etter, P. D. & Johnson, E. A. From promise to practice: pairing non-invasive sampling with genomics in conservation. PeerJ 3, e1106 (2015).

46. Danecek, P. et al. The variant call format and VCFtools. Bioinformatics 27, 2156–2158 (2011).

47. Jombart, T., Devillard, S. & Balloux, F. Discriminant analysis of principal components: a new method for the analysis of genetically structured populations. BMC Genet. 11, 94 (2010).

48. Alexander, D. H., Novembre, J. & Lange, K. Fast model-based estimation of ancestry in unrelated individuals. Genome Res. 19, 1655–1664 (2009).

49. Antao, T., Lopes, A., Lopes, R. J., Beja-Pereira, A. & Luikart, G. LOSITAN: A workbench to detect molecular adaptation based on a Fst -outlier method. BMC Bioinformatics 9, 1–5 (2008).

50. Luu, K., Bazin, E. & Blum, M. G. B. pcadapt: An R package to perform genome scans for selection based on principal component analysis. Mol. Ecol. Resour. 33, 67–77 (2016).

51. Lotterhos, K. E. & Whitlock, M. C. The relative power of genome scans to detect local adaptation depends on sampling design and statistical method. Mol. Ecol. 24, 1031–1046 (2015).

52. Huang, J. et al. Transcriptome Sequencing and Analysis of Leaf Tissue of *Avicennia marina* Using the Illumina Platform. PLoS One 9, e108785 (2014).

53. Lyu, H., Li, X., Guo, Z., He, Z. & Shi, S. *De novo* assembly and annotation of the *Avicennia officinalis* L. transcriptome. Mar. Genomics 38, 17–20 (2017).

54. Krishnamurthy, P. et al. Transcriptomics analysis of salt stress tolerance in the roots of the mangrove *Avicennia officinalis*. Nat. Sci. Reports 7, 1–19 (2017).

55. Park, S.-Y. et al. The Senescence-Induced Staygreen Protein Regulates Chlorophyll Degradation. Plant Cell 19, 1649–1664 (2007).

56. Myouga, F. et al. A Heterocomplex of Iron Superoxide Dismutases Defends Chloroplast Nucleoids against Oxidative Stress and Is Essential for Chloroplast Development in *Arabidopsis*. Plant Cell Online 20, 3148–3162 (2008).

57. Wang, S. et al. Chloroplast RNA-Binding Protein RBD1 Promotes Chilling Tolerance through 23S rRNA Processing in *Arabidopsis*. PLoS Genet. 12, 1–21 (2016).

58. Goulas, E. et al. The chloroplast lumen and stromal proteomes of *Arabidopsis thaliana* show differential sensitivity to short- and long-term exposure to low temperature. Plant J. 47, 720–734 (2006).

59. Kim, Y. O., Kim, J. S. & Kang, H. Cold-inducible zinc finger-containing glycine-rich RNA-binding protein contributes to the enhancement of freezing tolerance in *Arabidopsis thaliana*. Plant J. 42, 890–900 (2005).

60. Hartung, F., Suer, S. & Puchta, H. Two closely related RecQ helicases have antagonistic roles in homologous recombination and DNA repair in *Arabidopsis thaliana*. Proc. Natl. Acad. Sci. 104, 18836–18841 (2007).

61. Michaels, S. D. et al. AGL24 acts as a promoter of flowering in *Arabidopsis* and is positively regulated by vernalization. Plant J. 33, 867–874 (2003).

62. Jung, Y. J. et al. Universal Stress Protein Exhibits a Redox-Dependent Chaperone Function in *Arabidopsis* and Enhances Plant Tolerance to Heat Shock and Oxidative Stress. Front. Plant Sci. 6, 1–11 (2015).

63. Ogawa, T. et al. Functional Analysis of *Arabidopsis* Ethylene-Responsive Element Binding Protein Conferring Resistance to Bax. Plant Physiol. 138, 1436–1445 (2005).

64. Lin, R. C., Park, H. J. & Wang, H. Y. Role of *Arabidopsis* RAP2.4 in regulating light- and ethylene-mediated developmental processes and drought stress tolerance. Mol. Plant 1, 42–57 (2008).

65. Rae, L., Lao, N. T. & Kavanagh, T. A. Regulation of multiple aquaporin genes in *Arabidopsis* by a pair of recently duplicated DREB transcription factors. Planta 234, 429–444 (2011).

66. Alexandersson, E. et al. Whole gene family expression and drought stress regulation of aquaporins. Plant Mol. Biol. 59, 469–484 (2005).

67. Nishizawa, A., Yabuta, Y. & Shigeoka, S. Galactinol and Raffinose Constitute a Novel Function to Protect Plants from Oxidative Damage. Plant Physiol. 147, 1251–1263 (2008).

68. Zhang, A. et al. S-Type Anion Channels SLAC1 and SLAH3 Function as Essential Negative Regulators of Inward K^+^ Channels and Stomatal Opening in Arabidopsis. Plant Cell 28, 949–965 (2016).

69. Liepman, A. H. & Olsen, L. J. Alanine aminotransferase homologs catalyze the glutamate:glyoxylate aminotransferase reaction in peroxisomes of *Arabidopsis*. Plant Physiol. 131, 215–27 (2003).

70. Wolf, J. B. W., Lindell, J. & Backstrom, N. Speciation genetics: current status and evolving approaches. Philos. Trans. R. Soc. B Biol. Sci. 365, 1717–1733 (2010).

71. Gould, B. A., Chen, Y. & Lowry, D. B. Gene Regulatory Divergence Between Locally Adapted Ecotypes in Their Native Habitats. Mol. Ecol. 27, 4174–4188 (2018).

72. Ahrens, C. W. et al. The search for loci under selection: trends, biases and progress. Mol. Ecol. 27, 1342–1356 (2018).

73. Sakamoto, H., Araki, T., Meshi, T. & Iwabuchi, M. Expression of a subset of the *Arabidopsis* Cys2/His2-type zinc-finger protein gene family under water stress. Gene 248, 23–32 (2000).

74. Giuntoli, B. et al. A trihelix DNA binding protein counterbalances hypoxia-responsive transcriptional activation in *Arabidopsis*. PLoS Biol. 12, (2014).

75. Tohge, T. et al. Functional genomics by integrated analysis of metabolome and transcriptome of *Arabidopsis* plants over-expressing an MYB transcription factor. Plant J. 42, 218–235 (2005).

76. Bohne, A. V., Schwenkert, S., Grimm, B. & Nickelsen, J. Roles of Tetratricopeptide Repeat Proteins in Biogenesis of the Photosynthetic Apparatus. Int. Rev. Cell Mol. Biol. 324, 187–227 (2016).

77. Reef, R. & Lovelock, C. E. Regulation of water balance in Mangroves. Ann. Bot. 115, 385–395 (2015).

78. Menezes, M. P. M. de, Berger, U. & Mehlig, U. Mangrove vegetation in Amazonia: a review of studies from the coast of Pará and Maranhão States, north Brazil. Acta Amaz. 38, 403–420 (2008).

79. De Alvarenga, A. M. S. B., Botosso, P. C. & Soffiatti, P. Stem growth and phenology of three subtropical mangrove tree species. Brazilian J. Bot. 40, 907–914 (2017).

80. Kazan, K. & Lyons, R. The link between flowering time and stress tolerance. J. Exp. Bot. 67, 47–60 (2016).

81. Lovelock, C. E. et al. The vulnerability of Indo-Pacific mangrove forests to sea-level rise. Nature 526, 559–563 (2015).

82. Gall, H. Le et al. Cell Wall Metabolism in Response to Abiotic Stress. Plants 4, 112–166 (2015).

83. Kimura, M. et al. Identification of *Arabidopsis* Genes Regulated by High Light – Stress Using cDNA Microarray. Photochem. Photobiol. 77, 226–233 (2003).

84. Wang, D. et al. Genome-wide temporal-spatial gene expression profiling of drought responsiveness in rice. BMC Genomics 12, 149 (2011).

85. Moumeni, A. et al. Comparative analysis of root transcriptome profiles of two pairs of drought-tolerant and susceptible rice near-isogenic lines under different drought stress. BMC Plant Biol. 11, 174 (2011).

86. Wei, W. et al. Melatonin enhances plant growth and abiotic stress tolerance in soybean plants. J. Exp. Bot. 66, 695–707 (2015).

87. McRae, G. J. A Simple Procedure for Calculating Atmospheric Water Vapor Concentration. J. Air Pollut. Control Assoc. 30, 394–394 (1980).

88. Stuart, S. A., Choat, B., Martin, K. C., Holbrook, N. M. & Ball, M. C. The role of freezing in setting the latitudinal limits of mangrove forests. New Phytol. 173, 576–583 (2007).

89. Carlson, J. E., Holsinger, K. E. & Prunier, R. Plant responses to climate in the cape floristic region of South Africa: Evidence for adaptive differentiation in the proteaceae. Evolution (N. Y). 65, 108–124 (2011).

90. Pecot, S. D., Horsley, S. B., Battaglia, M. A. & Mitchell, R. J. The influence of canopy, sky condition, and solar angle on light quality in a longleaf pine woodland. Can. J. For. Res. 35, 1356–1366 (2005).

91. Arcus, V. L. et al. On the Temperature Dependence of Enzyme-Catalyzed Rates. Biochemistry 55, 1681–1688 (2016).

92. Cavanaugh, K. C. et al. Poleward expansion of mangroves is a threshold response to decreased frequency of extreme cold events. Proc. Natl. Acad. Sci. U. S. A. 111, 723–7 (2014).

93. Osland, M. J. et al. Climatic controls on the global distribution, abundance, and species richness of mangrove forests. Ecol. Monogr. 87, 341–359 (2017).

94. Colmer, T. D. & Flowers, T. J. Flooding tolerance in halophytes. New Phytol. 179, 964–974 (2008).

95. Schaeffer-Novelli, Y., Cintrón-Molero, G., Adaime, R. R. & de Camargo, T. M. Variability of mangrove ecosystems along the Brazilian coast. Estuaries 13, 204–218 (1990).

96. Godoy, M. D. P. & De Lacerda, L. D. Mangroves Response to Climate Change: A Review of Recent Findings on Mangrove Extension and Distribution. Ann. Brazilian Acad. Sci. 87, 651–667 (2015).

97. IPCC. Climate Change 2014: Synthesis Report. Contribution of Working Groups I, II and III to the Fifth Assessment Report of the Intergovernmental Panel on Climate Change. Cambridge University Press (IPCC, 2014).

98. Ellison, J. C. Vulnerability assessment of mangroves to climate change and sea-level rise impacts. Wetl. Ecol. Manag. 23, 115–137 (2015).

99. Heskel, M. A. et al. Convergence in the temperature response of leaf respiration across biomes and plant functional types. Proc. Natl. Acad. Sci. 113, 3832–3837 (2016).

100. Jordan, D. & Ogren, W. The CO2/O2 specificity of ribulose 1,5-bisphosphate carboxylase/oxygenase. Planta 161, 308–313 (1984).

101. Drake, B. G., Gonzàlez-Meler, M. A. & Long, S. P. More efficient plants: a consequence of rising atmospheric CO2? Annu. Rev. Plant Physiol. Plant Mol. Biol. 48, 609–639 (1997).

102. Saenger, P. & Moverly, J. Vegetative phenology along the Queensland coastline. Proc. Ecol. Soc. Aust. 13, 257–265 (1985).

103. Doughty, C. E. et al. Drought impact on forest carbon dynamics and fluxes in Amazonia. Nature 519, 78–82 (2015).

104. Rodell, M. et al. Emerging trends in global freshwater availability. Nature 557, 651–659 (2018).

105. Osland, M. J. et al. Climate and plant control on soil organic matter in coastal wetlands. Glob. Chang. Biol. (2018). doi:10.1111/gcb.14376

106. Bulleri, F. et al. Harnessing positive species interactions as a tool against climate-driven loss of coastal biodiversity. Plos Biol. 16, (2018).

107. Moritz, C. Defining ‘Evolutionarily Significant Units’ for conservation. Trends Ecol. Evol. 9, 373–375 (1994).

108. Ikeda, D. H. et al. Genetically informed ecological niche models improve climate change predictions. Glob. Chang. Biol. 23, 164–176 (2017).

109. Holliday, J. A. et al. Advances in ecological genomics in forest trees and applications to genetic resources conservation and breeding. Mol. Ecol. 706–717 (2017). doi:10.1111/mec.13963

110. Vestbo, S., Obst, M., Quevedo Fernandez, F. J., Intanai, I. & Funch, P. Present and Potential Future Distributions of Asian Horseshoe Crabs Determine Areas for Conservation. Front. Mar. Sci. 5, (2018).

111. Hijmans, R. J., Cameron, S. E., Parra, J. L., Jones, P. G. & Jarvis, A. Very high resolution interpolated climate surfaces for global land areas. Int. J. Climatol. 25, 1965–1978 (2005).

112. Fick, S. E. & Hijmans, R. J. WorldClim 2: new 1-km spatial resolution climate surfaces for global land areas. Int. J. Climatol. 4315, 4302–4315 (2017).

113. Sbrocco, E. J. & Barber, P. H. MARSPEC: ocean climate layers for marine spatial ecology. Ecology 94, 979–979 (2013).

114. Forsythe, W. C., Rykiel, E. J., Stahl, R. S., Wu, H. i. & Schoolfield, R. M. A model comparison for daylength as a function of latitude and day of year. Ecol. Modell. 80, 87–95 (1995).

