## Supplementary for "Local adaptation of a dominant coastal tree to freshwater availability and solar radiation suggested by genomic and ecophysiological approaches"

### Supplementary information

**Article title:** Local adaptation to water-deficit and light limitation suggested from genomic and ecophysiological approaches in a dominant coastal tree

**Authors:** Mariana Vargas Cruz, Gustavo Maruyama Mori, Caroline Signori Müller, Carla Cristina da Silva, Dong-Ha Oh, Maheshi Dassanayake, Maria Imaculada Zucchi, Rafael Silva Oliveira and Anete Pereira de Souza

The following Supplementary information is available for this article:

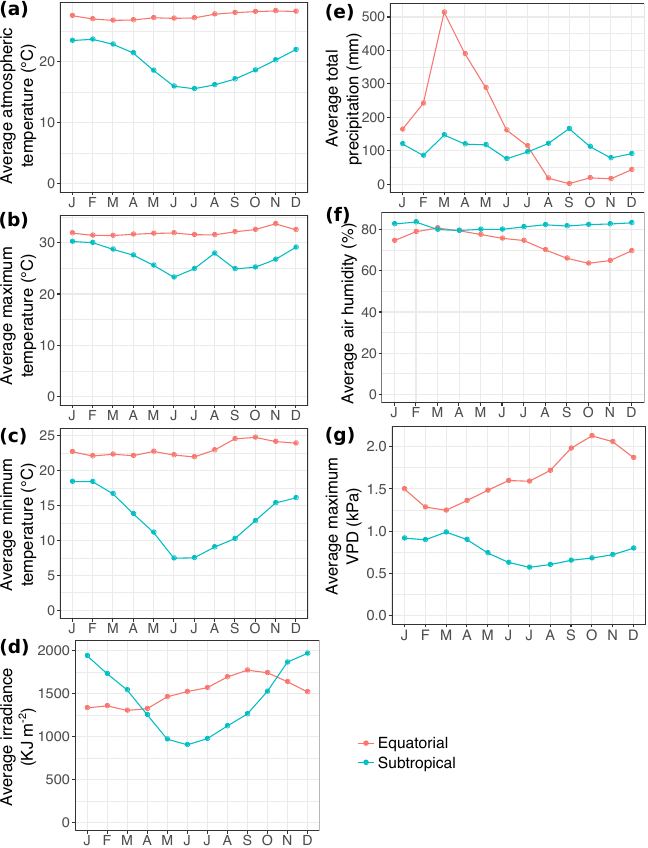

**Supplementary Fig. S1.** Climate characterisation of equatorial and subtropical sites. Lines represent the monthly means of climate variables (2008-2017). Red: equatorial site (Tracuateua automatic station); blue: subtropical site (Santa Marta automatic station). Source: Brazilian National Institute of Meteorology (INMET). (a) Average temperature; (b) average maximum temperature; (c) average minimum temperature; (d) average solar irradiance; (e) total precipitation; (f) average air humidity; (g) average air vapour pressure deficit (VPD).

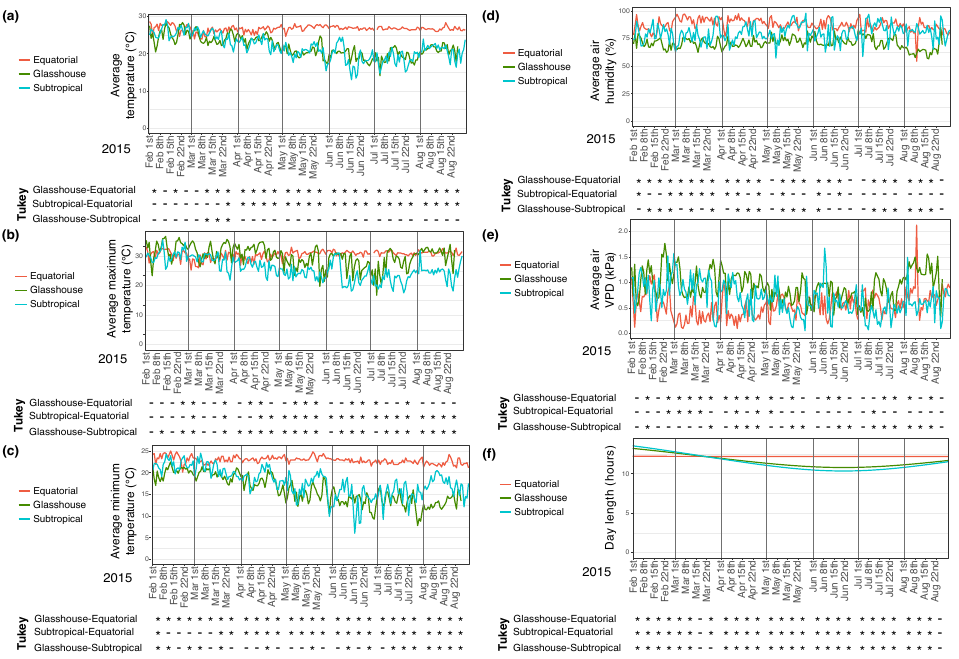

**Supplementary Fig. S2.** Environmental data in the glasshouse and at the source sites of propagules used in the common garden experiment. Green line: glasshouse; red line: equatorial site (Tracuateua automatic station, source: Brazilian National Institute of Meteorology (INMET)); blue line: subtropical site (Santa Marta automatic station, source: INMET). Multiple comparisons of periodic data of each environmental variable measured during the experiment were performed using ANOVA and post-hoc Tukey HSD tests with a significance threshold of 0.05. * Represent significant differences and hyphens (-) represent an absence of a significant difference. (a) Average temperature; (b) average maximum temperature; (c) average minimum temperature; (d) average air humidity; (e) average air vapour pressure deficit (VPD).

**Supplementary Fig. S3.** *Avicennia schaueriana* from equatorial and subtropical sites grown in common garden. (a) Geographical distribution of the species (green shaded zone) and location of the equatorial (red dot) and the subtropical (blue dot) sampling sites of propagules used in the experiment conducted in a green house (black square); (b) propagules germinating in trays filled with mangrove soil; (c-d) seedlings from the equatorial and from the subtropical sampling sites on the 80th day of the experiment; (e-f) seedlings from the equatorial and subtropical sampling sites on the 225th day of the experiment; (g) seven-months-old *Avicennia schaueriana* from the equatorial sampling site presenting flower buds and flowers; (h-i) micrographs of stem transverse sections from pith to the outermost growth section of one representative seedling from the equatorial and subtropical sampling sites (magnification = 10x); stem sections were stained with Astra Blue and Safranin O; V: xylem vessels; Ph: phloem; CCC: vascular cambium.

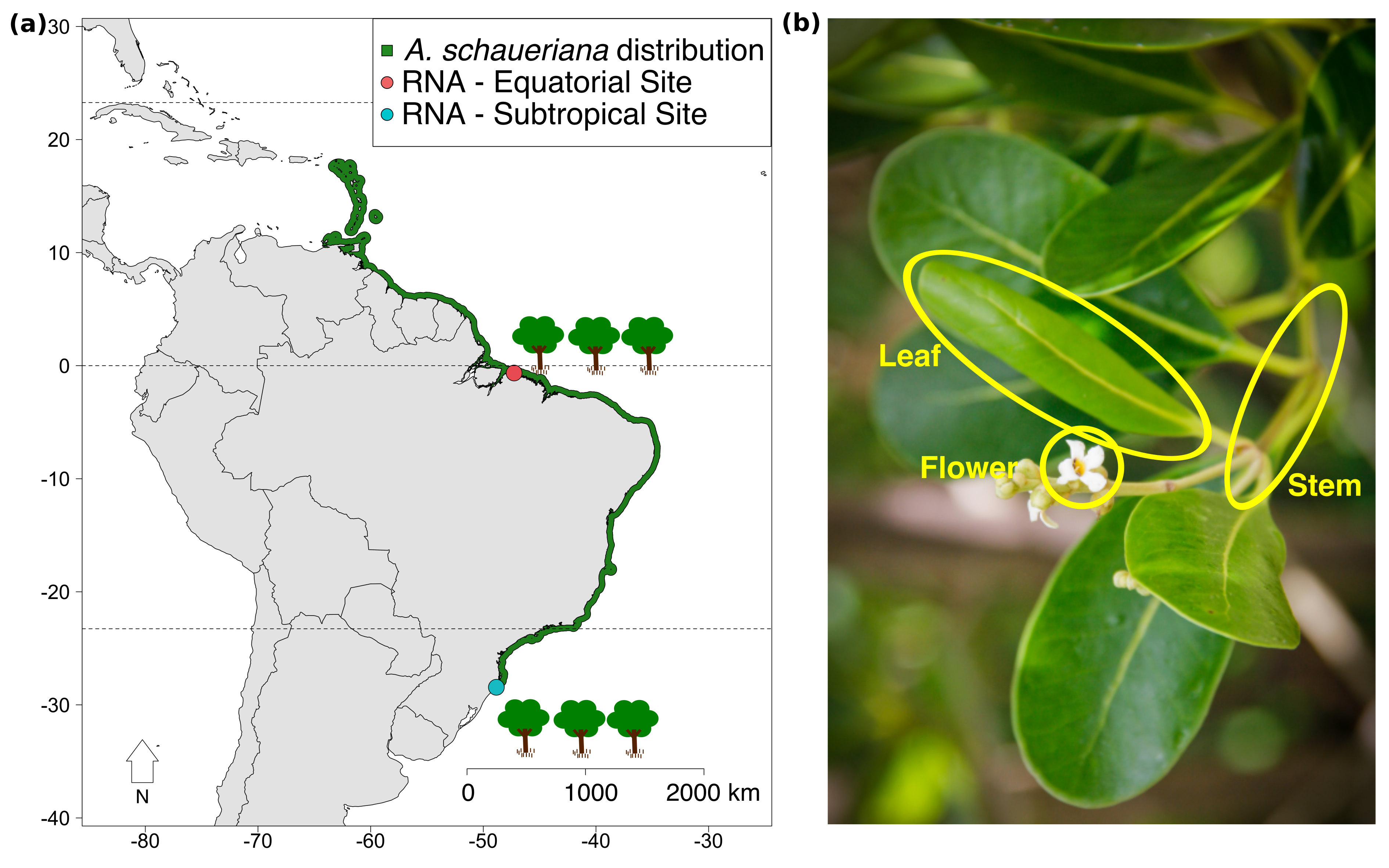

**Supplementary Fig. S4.** Sampling of *Avicennia schaueriana* plant material used for RNA-Sequencing. (a) Geographical distribution of the species (green-shaded area) and the equatorial (red dot) and subtropical (blue dot) sites; (b) plant organs sampled in the field for RNA extraction and sequencing (bottom).

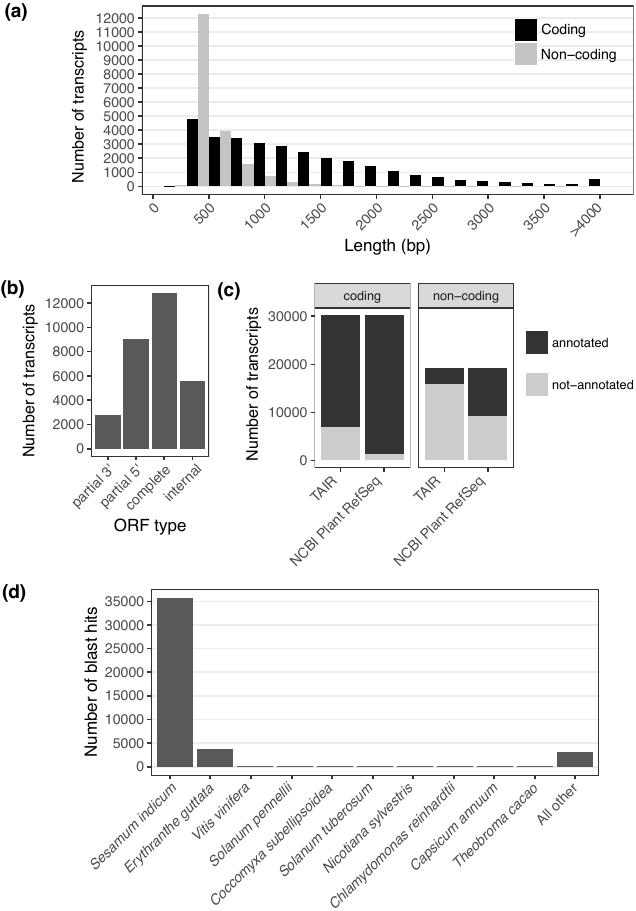

**Supplementary Fig. S5.** Characterisation of the de novo assembly of the *Avicennia schaueriana* reference transcriptome. (a) Histogram of the length of putative protein-coding and putative non-coding transcripts; (b) classification of all predicted open reading frames (ORF) in the reference transcriptome: complete, partial 3’ (missing stop codon), partial 5’ (missing start codon), internal (missing both start and stop codons); (c) annotation of transcripts via blastx using The *Arabidopsis* Information Resource (TAIR) protein database and The National Center for Biotechnology Information (NCBI) RefSeq Plant protein database; (d) blastx top-hit species distribution of transcripts annotations using NCBI RefSeq Plant protein database as reference.

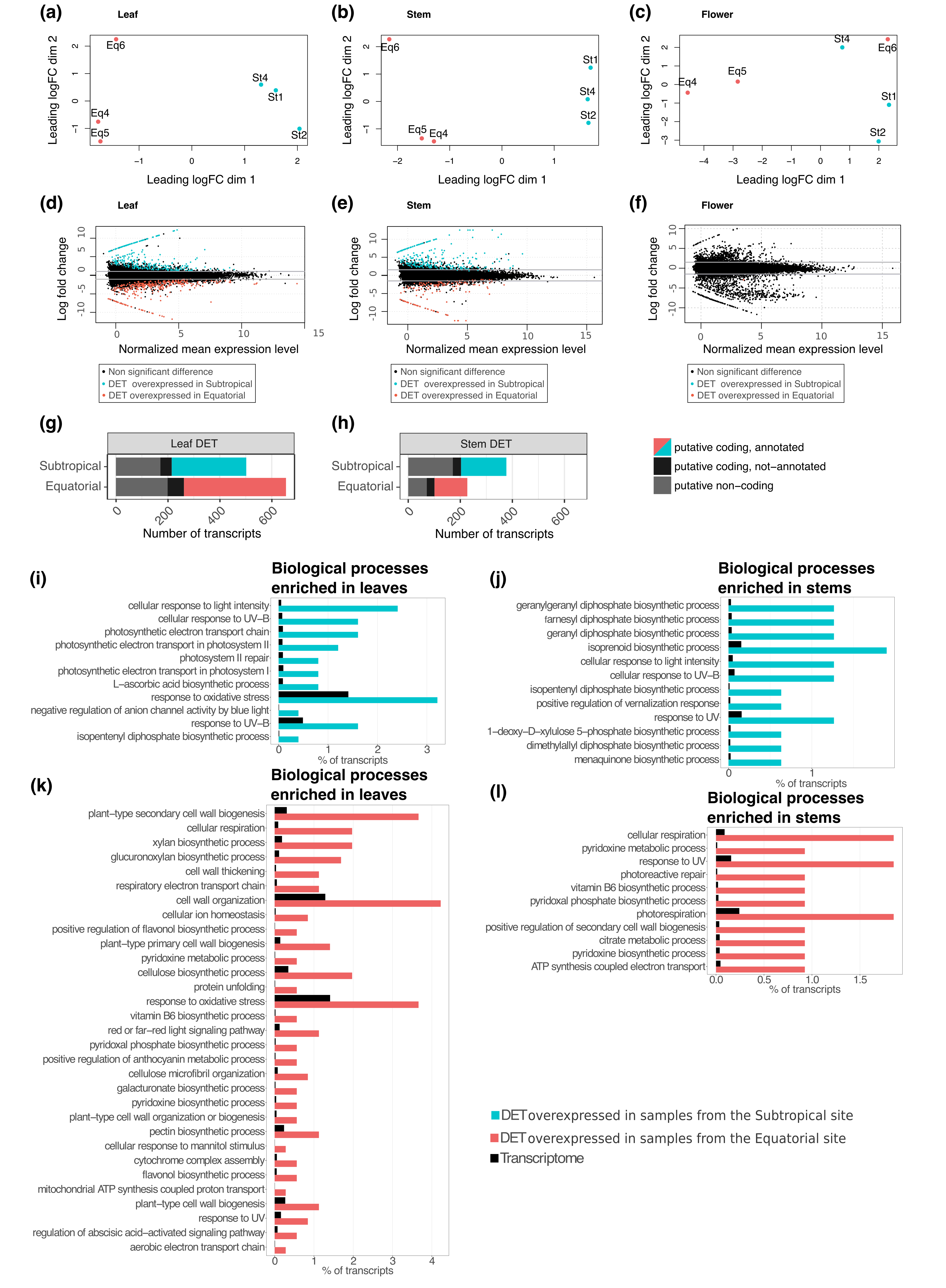

**Supplementary Fig. S6.** (Previous page) Detection of differentially expressed transcripts (DET) and Gene Ontology (GO) enrichment analysis. (a-c) Two-dimensional scatterplot of distances among transcript expression levels in leaf, stem and flower samples (left to right) of *Avicennia schaueriana* trees under field conditions; points represent individual samples, and distances are approximately the log2-transformed fold changes between samples; red points represent samples from the equatorial site and blue points represent samples from the subtropical site. (d-f) MA-plots of transcript expression in leaves, stems and flowers; red points represent DET overexpressed in equatorial samples; blue points represent DET overexpressed in subtropical samples; black points represent non-differentially expressed transcripts; grey horizontal lines represent ± 1 log fold-change threshold. (g-h) Proportion of DET used in the GO enrichment analysis represented in blue and red bars. Annotation of putative coding genes was obtained using the blastx algorithm with *Arabidopsis thaliana* proteins as a reference; grey bars represent putative non-coding transcripts that are differentially expressed, and black bars represent putative coding transcripts that are differentially expressed but not annotated. (i-l) Examples of Gene Ontology (GO) terms enriched among overexpressed transcripts (DET) in subtropical leaf samples; in equatorial leaf samples; in subtropical stem samples and in equatorial stem samples. GO terms are ordered from the top down based on the P- value of enrichment (low to high, all < 0.05). Coloured bars represent the percentage of all DET that belong to each of the listed categories. Black bars represent the percentage of all transcripts in the reference transcriptome belonging to each category. Plant material of adult trees was sampled under field conditions, during the end of winter in the subtropical site and in the beginning of the dry season in the equatorial site. The complete lists of enriched GO terms are presented in Supplementary information Tables S7-S10.

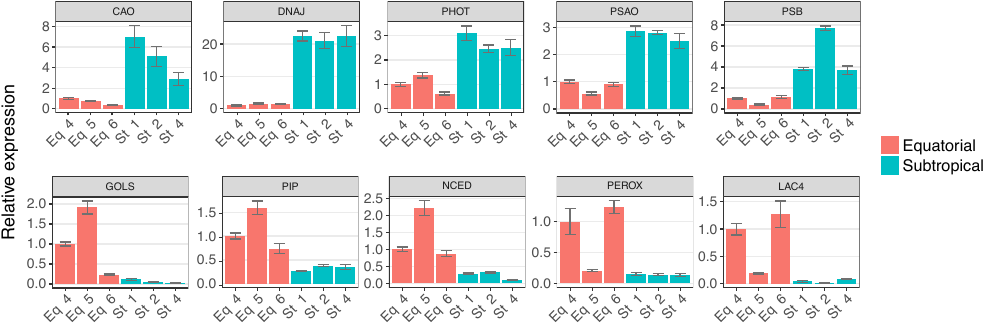

**Supplementary Fig. S7.** Validation of RNA-Sequencing-based differential expression of transcripts by reverse transcription real time PCR (qRT-PCR). Each bar represents the relative expression of one sequenced sample. Error bars represent the standard deviation of the mean from three technical replicates of the same individual sample. Red bars represent samples from the equatorial sampling site and blue bars, samples from the subtropical sampling site. Non-parametric unpaired Mann-Whitney Wilcoxon U-tests were used to compare the distributions. The differential expression detected *in silico* was confirmed *in vitro* for all putative genes, at a significance threshold of 0.05. Putative genes selected for the validation of the RNA-sequencing data and primers sequences are described in Supplementary information Table S12.

**Supplementary Table S1.** Characterisation of field conditions at the time of plant material collection and storage in RNA-stabilising solution for subsequent sequencing.

| Sample  ID | Sampling site  (Climate^†^) | GPS Point | Date (d/m) | Time (h:m) | TL (m) | Temp (°C)^‡^ | RH (%)^‡^ | VPD (kPa)^§^ | SI (KJ/m^2^)^‡^ |
| --- | --- | --- | --- | --- | --- | --- | --- | --- | --- |
| St1 | Subtropical  (Cfa) | 28.48 S; 48.88 W | 23/08 | 15:15 | 0.5 | 19 | 84 | 0.351 | 2033.89 |
| St2 | Subtropical  (Cfa) | 28.36 S; 48.80 W | 24/08 | 10:35 | 0.5 | 18.3 | 86 | 0.294 | 1804.15 |
| St4 | Subtropical  (Cfa) | 28.48 S; 48.83 W | 25/08 | 13:11 | 0.5 | 20.9 | 82 | 0.445 | 1863.14 |
| Eq4 | Equatorial  (Am) | 00.64 S; 47.26 W | 28/07 | 11:57 | 2.7 | 28.5 | 49 | 1.983 | 1940.73 |
| Eq5 | Equatorial  (Am) | 00.65 S; 47.26 W | 29/07 | 12:22 | 2.7 | 28.3 | 51 | 1.883 | 2724.54 |
| Eq6 | Equatorial  (Am) | 00.64 S; 47.26 W | 29/07 | 16:00 | 0.7 | 30.1 | 47 | 2.260 | 1433.68 |

TL: tidal level; Temp: atmospheric air temperature; RH: atmospheric relative humidity; VPD: air vapour pressure deficit; SI: solar irradiance.

^†^Köppen-Geiger Climate classification according to Alvares et al.^1^.

^‡^Source: Brazilian National Institute of Meteorology (INMET).

^§^Estimated according to McRae^2^.

**Supplementary Table S2.** Sampling sites for plant material used in nextRAD sequencing.

| Site ID | Number of samples | Cluster^†^ | Locality (City, State) | Geographic Coordinate |
| --- | --- | --- | --- | --- |
| PAR | 9 | N | Bragança, Pará | 0° 49' 12" S, 46° 36' 56" W |
| ALC | 8 | N | Alcântara, Maranhão | 2° 24' 37" S, 44° 24' 22" W |
| PRC | 9 | N | Paracuru, Ceará | 3° 24' 47" S, 39° 3' 23" W |
| VER | 9 | S | Vera Cruz, Bahia | 12° 59' 1" S, 38° 41' 5" W |
| GPM | 9 | S | Guapimirim, Rio de Janeiro | 22° 42' 5" S, 43° 0' 26" W |
| UBA | 9 | S | Ubatuba, São Paulo | 23° 29' 22" S, 45° 9' 52" W |
| CNN | 8 | S | Cananéia, São Paulo | 25° 1' 12" S, 47° 55' 5" W |
| PPR | 8 | S | Pontal do Paraná, Paraná | 25° 34' 30" S, 48° 21' 9" W |
| FLN | 7 | S | Florianópolis, Santa Catarina | 27° 34' 37" S, 48° 31' 8" W |
| LGN | 3 | S | Laguna, Santa Catarina | 28° 29' 44'' S, 48° 53' 28 W |

^†^Cluster classification based on the structure of diversity in single nucleotide polymorphisms (SNP) in DNA sequences. N: North from the Northeast Extremity of South America; S: South from the Northeast Extremity of South America.

**Supplementary Table S3.** Morphological traits analysed in *Avicennia schaueriana* saplings from Equatorial and Subtropical sites, grown under the same environmental conditions during seven months in a glasshouse.

| Variable |  | Equatorial | | |  | Subtropical | | |  | P-value^†^ |
| --- | --- | --- | --- | --- | --- | --- | --- | --- | --- | --- |
|  |  | mean |  | sd |  | mean |  | sd |  |  |
| Leaves dry weight (g) |  | 3.78 |  | (1.05) |  | 7.41 |  | (1.73) |  | 4.18e-07 (T) |
| Stems dry weight (g) |  | 4.16 |  | (1.67) |  | 5.33 |  | (1.51) |  | 0.05332 (T) |
| Roots dry weight (g) |  | 4.97 |  | (1.57) |  | 8.24 |  | (2.36) |  | 1.57e-04 (T) |
| Leaf dry mass ratio (%) (Relative to total) |  | 29.67 |  | (2.85) |  | 35.53 |  | (3.00) |  | 7.17e-06 (T) |
| Stem dry mass ratio (%) (Relative to total) |  | 31.47 |  | (3.56) |  | 25.33 |  | (2.66) |  | 1.38e-05 (T) |
| Root dry mass ratio (%) (Relative to total) |  | 38.73 |  | (3.47) |  | 39.20 |  | (3.05) |  | 0.699 (T) |
| Individual leaf area (cm²) |  | 3.32 |  | (2.24) |  | 6.04 |  | (4.24) |  | 2.20e-16 (W) |
| Total leaf area (cm²) |  | 363.14 |  | (119.55) |  | 507.28 |  | (135.23) |  | 0.120 (T) |
| Specific Leaf Area (cm²/g) |  | 116.38 |  | (28.25) |  | 101.81 |  | (18.56) |  | 0.3044 (T) |
| Leaf lamina angle (degrees) |  | 48.85 |  | (6.38) |  | 69.21 |  | (10.35) |  | 1.02e-06 (T) |
| Vessel lumen area (μm²) |  | 180.18 |  | (72.03) |  | 280.37 |  | (119.49) |  | < 2.20e-16 (W) |
| Vessel lumen diameter (μm) |  | 14.87 |  | (2.89) |  | 18.50 |  | (3.83) |  | < 2.20e-16 (W) |
| Vessel density in the xylem (number of vessels per mm²) |  | 162.40 |  | (17.61) |  | 185.43 |  | (44.28) |  | 0.236 (T) |
| Vessel lumen area in sapwood (%) |  | 2.20 |  | (0.55) |  | 3.91 |  | (1.14) |  | 0.1055 (T) |
| Vessel grouping index |  | 2.90 |  | (0.29) |  | 2.36 |  | (0.15) |  | 0.081 (W) |

^†^Unpaired two-group test. Mann-Whitney-Wilcoxon (W) or Student’s T-test (T).

**Supplementary Table S4.** Transcriptome quality parameters assessed in this work.

| Reads mapping to contiguous sequences in the transcriptome^†^ | Mapped to a single contig | 91.91% |
| --- | --- | --- |
|  | Mapped to more than one contig | 1.16% |
|  | Not mapped to the transcriptome | 6.93% |
| Plant universal single-copy orthologs present in the transcriptome^‡^ | Complete orthologous sequences present in single copy in the transcriptome | 64.12% |
|  | Complete orthologous sequences present in more than one copy in the transcriptome | 21.65% |
|  | Fragmented orthologous sequences present in the transcriptome | 6.07% |
|  | Absent orthologous sequences in the transcriptome | 8.16% |

^†^Analysis performed using Bowtie^3^

^‡^BUSCO analysis^4^.

**Supplementary Table S5.** Transcriptome size characterisation.

|  |  | All contigs |  | Putative coding contigs |  | Putative non-coding contigs |
| --- | --- | --- | --- | --- | --- | --- |
| Number of contigs |  | 49,490 |  | 30,227 |  | 19,263 |
| Total size (bp) |  | 49,860,064 |  | 40,048,718 |  | 9,811,346 |
| Shortest contig (bp) |  | 282 |  | 298 |  | 282 |
| Longest contig (bp) |  | 11,672 |  | 11,672 |  | 3,760 |
| Number of contigs < 1Kb |  | 31,787 |  | 13,406 |  | 18,373 |
| Number of contigs ≥ 1Kb |  | 17,703 |  | 16,822 |  | 881 |
| Average contig size (bp) |  | 1,007 |  | 1,324 |  | 509 |
| Median contig size (bp) |  | 702 |  | 1,116 |  | 427 |
| Contig N50 (bp) |  | 1,428 |  | 1,708 |  | 515 |
| Average ORF size (Aac) |  | NA |  | 332 |  | NA |
| Median ORF size (Aac) |  | NA |  | 253 |  | NA |
| Longest ORF (Aac) |  | NA |  | 3,755 |  | NA |
| Shortest ORF (Aac) |  | NA |  | 100 |  | NA |

**Supplementary Table S6.** Putative orthologous expressed sequences between *Avicennia schaueriana* transcripts obtained from flowers, stems and leaves and publicly available transcriptomes from other *Avicennia* species.

|  | *Avicennia marina* |  | *Avicennia officinalis* |  | *Avicennia officinalis* |
| --- | --- | --- | --- | --- | --- |
| Plant material | Leaves |  | Leaves |  | Roots |
| NCBI or DDBJ/EMBL/GenBank accession number | GBIO01000000 |  | GFLY01000000 |  | GSE73807 |
| Number of transcripts | 89,833 |  | 38,756 |  | 121,929 |
| Total number of putative orthologs | 27,658 |  | 18,325 |  | 13,273 |
| Number of putative protein coding orthologs | 21,313 |  | 17,663 |  | 9,993 |
| Reference | Huang *et al.* (2014)^5^ |  | Lyu *et al.* (2017)^6^ |  | Krishnamurthy *et al.* (2017)^7^ |

**Supplementary Table S7.** Enriched Gene Ontology (GO) terms among differentially expressed transcripts (DET) that presented higher expression in subtropical stem samples than in equatorial stem samples. (Too large to be included in this file. Submitted as separate file).

**Supplementary Table S8.** Enriched Gene Ontology (GO) terms among differentially expressed transcripts (DET) that presented higher expression in equatorial stem samples than in subtropical stem samples. (Too large to be included in this file. Submitted as separate file).

**Supplementary Table S9.** Enriched Gene Ontology (GO) terms among differentially expressed transcripts (DET) that presented higher expression in subtropical leaf samples than in equatorial leaf samples. (Too large to be included in this file. Submitted as separate file).

**Supplementary Table S10.** Enriched Gene Ontology (GO) terms among differentially expressed transcripts (DET) that presented higher expression in equatorial leaf samples than in subtropical leaf samples. (Too large to be included in this file. Submitted as separate file).

**Supplementary Table S11.** Examples of differentially expressed transcripts (DET) of *Avicennia schaueriana* associated to the response to contrasting climate conditions between equatorial and subtropical sites. (Too large to be included in this file. Submitted as separate file).

**Supplementary Table S12.** Oligonucleotides used in qRT-PCR reactions in this work. (Too large to be included in this file. Submitted as separate file).

**Supplementary Table S13.** Single Nucleotide Polymorphic loci (SNP) with signature of selection present in putative protein-coding regions of the genome of *Avicennia schaueriana*. (Too large to be included in this file. Submitted as separate file).

**Supplementary Methods**

***Propagules and RNA sampling sites characterisation***

Here we complement the characterisation of the sites where leaves, stems and flowers of adult individuals of *Avicennia schaueriana* were collected for RNA-sequencing and propagules were sampled for a common garden experiment.

The equatorial site is located in the Amazon mangrove coast, one of the world’s largest macrotidal mangrove forests^8,9^ (0°42' S 47°14' W and 0°59' S 46°35' W). Forests reach 20 m in height and are dominated by *Rhizophora mangle* and *R. racemosa,* at lower elevations, and by *Avicennia germinans* at higher elevations^10,11^. *Avicennia schaueriana* and *Laguncularia racemosa* are frequently found at lower densities at the edges of forests in the equatorial coast, such that the former occurs especially close to sandy beaches, whereas the latter occurs in brackish water^10,11^. The climate at the equatorial site is classified as tropical monsoon (Am) in the Köppen-Geiger classification system. A decreasing rainfall gradient is observed from this sampling site towards the northeast extremity of South America (NEESA), where the climate is semi-arid of low latitude and altitude (BSh)^1^. The mean annual temperature at this location is above 27 °C and exhibits great fluctuations in precipitation throughout the year. The dry season lasts for approximately five months, from August to December, with a precipitation lower than 50 mm, decreasing to 9 mm in October, which is the driest month. The wet period duration is approximately seven months, from January to July, with precipitation exceeding 300 mm, and with an average precipitation of 600 mm in the wettest month, March.

The subtropical sampling site is located at the southernmost margin of mangrove forests on the Atlantic coast of South America (28°28’ S 48°51’ W)^12^. Only two species of mangrove trees are found at this site, namely, *A. schaueriana* and *L. racemosa*. Despite being found at the edge of the species distribution, *A. schaueriana* individuals present a tall form, reaching up to 10 m in height, which suggests that their distribution may not have been limited by climatic factors^12^. The climate in this locality is classified as temperate oceanic with hot summer, without a dry season (Cfa) in the Köppen-Geiger classification system, with increasing temperatures and tidal amplitudes northwards, where climate is tropical with a dry season (Aw or As)^1^, in the NEESA region. The tidal regime in this subtropical site is classified as microtidal^12^ and the precipitation is well distributed and exceeds 90 mm throughout the year. The wettest month is February, with an average precipitation of 160 mm. The annual mean temperature is 19.7 °C, the mean temperature in the coldest month is 15.7 °C and the mean annual thermal amplitude is 8 °C.

***RNA-sequencing data verification through qRT-PCR***

To confirm the results obtained in the differential expression analyses of RNA-Seq data, we selected a set of DET detected between leaf samples of trees at the equatorial and subtropical sites and designed primers for reverse transcription real-time PCR (qRT-PCR) validation using the Primer3Plus software^13^. To design primers to be used as internal controls, we selected transcripts from the RNA-Seq data that showed a fairly stable expression across leaf samples and were similar to recognised housekeeping genes in genomes of model-plants Supplementary information Table 10). RNA samples were treated with the DNA-free DNA Removal Kit (Thermo Fisher Scientific, Waltham, MA, USA), followed by complementary DNA strand (cDNA) synthesis using the commercial iScript cDNA Synthesis kit (Bio-Rad Laboratories Inc., Hercules, CA, USA). Primers annealing specificity, amplification efficiency and the stability of expression of the putative housekeeping transcripts were all confirmed before performing qRT-PCR reactions. Each reaction was assessed using three technical replicates, and expression values were estimated with the 2^-ΔΔCt^ method^14^. The Mann-Whitney-Wilcoxon non-paired statistical test was performed with a significance threshold of 5%, for the comparison of the distributions of relative expression levels. All reactions were performed using the iTaq Universal SYBR Green Supermix (Bio-Rad Laboratories Inc., Hercules, CA, USA) with the CFX384 Real-Time PCR Detection System (Bio-Rad Laboratories Inc., Hercules, CA, USA), following the manufacturer instructions.

To better characterise the climate from the equatorial and subtropical sampling sites we have downloaded temperature, relative humidity, rainfall and solar irradiance datasets (2008-2017) from the INMET database (the Meteorological Institute of Brazil) (Supplementary information Fig. 1) and the environmental data from BioClim^15^ and MARSPEC^16^.

**Supplementary Note**

***Validation of RNA-Seq data through qRT-PCR***

To confirm the computational analyses of the RNA-Seq data, we selected a set of transcripts with differential and approximately stable expression across leaf samples and designed primers for use in reverse transcription real-time PCR (qRT-PCR) reactions. Two putative housekeeping genes, similar to *Arabidopsis thaliana* *UBIQUITIN-CONJUGATING ENZYME 11* and *UBIQUITIN-SPECIFIC PROTEASE 12*, were selected as reference in qRT-PCR validation of the RNA-Seq data, from leaves sampled from adult trees under equatorial and subtropical field conditions. The stability of the expression of the putative housekeeping genes was confirmed, showing a coefficient of variation (CV) of 0.0456 and mean variation (M) of 0.1316. Amplification efficiency tests for all primers were performed before examining the relative expression qRT-PCR reactions. Only primers with 90-110% efficiency and R^2^ > 0.99 were used in the subsequent steps. The specificity was confirmed by the analysis of the dissociation curve over a temperature gradient from 65 °C - 95 °C, with an increment of 0.5 °C every 5 seconds. Ten differentially expressed transcripts (DET), detected using read-mapping counts and the results obtained from the EdgeR^17^ analysis were selected as targets in the qRT-PCR reactions (Supplementary Table 10). Their relative expression, estimated using the 2^-ΔΔCt^ method^14^, confirmed the results obtained in computational analyses of the RNA-Seq data. An absence of a difference in the distribution of expression levels between equatorial and subtropical leaf samples was rejected for all selected targets using the non-parametric unpaired Mann-Whitney-Wilcoxon U statistic with a significance threshold of 0.05 (Supplementary information Fig. S8).

**References**

1.	Alvares, C. A., Stape, J. L., Sentelhas, P. C., Gonçalves, J. L. de M. & Sparovek, G. Köppen’s climate classification map for Brazil. *Meteorol. Zeitschrift* **22,** 711–728 (2013).

2.	McRae, G. J. A Simple Procedure for Calculating Atmospheric Water Vapor Concentration. *J. Air Pollut. Control Assoc.* **30,** 394–394 (1980).

3.	Langmead, B., Trapnell, C., Pop, M. & Salzberg, S. Ultrafast and memory-efficient alignment of short DNA sequences to the human genome. *Genome Biol.* **10,** R25 (2009).

4.	Simão, F. A., Waterhouse, R. M., Ioannidis, P., Kriventseva, E. V. & Zdobnov, E. M. BUSCO: Assessing genome assembly and annotation completeness with single-copy orthologs. *Bioinformatics* **31,** 3210–3212 (2015).

5.	Huang, J. *et al.* Transcriptome Sequencing and Analysis of Leaf Tissue of Avicennia marina Using the Illumina Platform. *PLoS One* **9,** e108785 (2014).

6.	Lyu, H., Li, X., Guo, Z., He, Z. & Shi, S. De novo assembly and annotation of the Avicennia officinalis L. transcriptome. *Mar. Genomics* 2–5 (2017). doi:10.1016/j.margen.2017.07.002

7.	Krishnamurthy, P. *et al.* Transcriptomics analysis of salt stress tolerance in the roots of the mangrove Avicennia officinalis. *Nat. Sci. Reports* **7,** 1–19 (2017).

8.	Souza-Filho, P. W. M. *et al.* Holocene Coastal Evolution and Facies Model of the Bragança Macrotidal Flat on the Amazon Mangrove Coast, Northern Brazil. *J. Coast. Res.* 306–310 (2006).

9.	Kjerfve, B. *et al.* Chapter Twenty Morphodynamics of muddy environments along the Atlantic coasts of North and South America. *Proc. Mar. Sci.* **4,** 479–532 (2002).

10.	Schaeffer-Novelli, Y., Cintrón-Molero, G., Adaime, R. R. & de Camargo, T. M. Variability of mangrove ecosystems along the Brazilian coast. *Estuaries* **13,** 204–218 (1990).

11.	Menezes, M. P. M. de, Berger, U. & Mehlig, U. Mangrove vegetation in Amazonia: a review of studies from the coast of Pará and Maranhão States, north Brazil. *Acta Amaz.* **38,** 403–420 (2008).

12.	Soares, M. L. G., Estrada, G. C. D., Fernandez, V. & Tognella, M. M. P. Southern limit of the Western South Atlantic mangroves: Assessment of the potential effects of global warming from a biogeographical perspective. *Estuar. Coast. Shelf Sci.* **101,** 44–53 (2012).

13.	Untergasser, A. *et al.* Primer3-new capabilities and interfaces. *Nucleic Acids Res.* **40,** 1–12 (2012).

14.	Livak, K. J. & Schmittgen, T. D. Analysis of relative gene expression data using real-time quantitative PCR and the 2(-Delta Delta C(T)) Method. *Methods* **25,** 402–8 (2001).

15.	Hijmans, R. J., Cameron, S. E., Parra, J. L., Jones, P. G. & Jarvis, A. Very high resolution interpolated climate surfaces for global land areas. *Int. J. Climatol.* **25,** 1965–1978 (2005).

16.	Sbrocco, E. J. & Barber, P. H. MARSPEC: ocean climate layers for marine spatial ecology. *Ecology* **94,** 979–979 (2013).

17.	Robinson, M. D., McCarthy, D. J. & Smyth, G. K. edgeR: a Bioconductor package for differential expression analysis of digital gene expression data. *Bioinformatics* **26,** 139–140 (2010).
